# α-Endosulfine regulates amyloid β 42 via the modulation of neprilysin activity

**DOI:** 10.1101/2020.10.07.329318

**Authors:** Naoto Watamura, Naomasa Kakiya, Per Nilsson, Satoshi Tsubuki, Naoko Kamano, Mika Takahashi, Shoko Hashimoto, Hiroki Sasaguri, Takashi Saito, Takaomi C. Saido

## Abstract

The neuropeptide somatostatin (SST) regulates amyloid β peptide (Aβ) catabolism by enhancing neprilysin (NEP)-catalyzed proteolytic degradation. However, the mechanism by which SST regulates NEP activity remains unclear. Here we report the identification by differential proteomics of α-endosulfine (ENSA), an endogenous ligand of the ATP-sensitive potassium (K_ATP_) channel, as a negative regulator of NEP activity downstream of SST signaling. Genetic deficiency of ENSA resulted in enhanced NEP activity and decreased Aβ deposition in the brains of wild-type and Alzheimer’s disease (AD) model mice. Pharmacological intervention to increase the probability of K_ATP_ channel opening reduced Aβ deposition in AD model mice. Our findings provide new insights into possible mechanisms to prevent AD.

## Introduction

Accumulation of amyloid β peptide (Aβ) in the brain is a pathological hallmark of Alzheimer’s disease (AD). To this end, the identification of pathogenic mutations in the *APP, PSEN1* and *PSEN2* genes supports the amyloid cascade hypothesis underlying the etiology of AD^1^, and verify that these mutations cause early-onset AD due to the abnormal production and accumulation of toxic Aβ species such as Aβ_42_ and Aβ_43_^2,3^. In contrast, the exact causes of Aβ deposition in sporadic AD cases remain unclear, although some genetic risk factors related to Aβ metabolism have been identified^4^.

Neprilysin (NEP; neutral endopeptidase 24.11) is a major physiological Aβ-degrading enzyme^5,6^, the expression of which in the brain declines with aging and in early stages of AD progression^7,8^. Identification of the mechanism(s) that regulate NEP activity should contribute to the development of ways to prevent AD. We previously showed that somatostatin (SST), a neuropeptide known as a somatotropin-release inhibiting hormone^9^, regulates Aβ_42_ levels in the brain via the upregulation of NEP activity^10^. In addition, we discovered that, of the five SST receptor (SSTR1-5) subtypes, SSTR1 and SSTR4 redundantly regulate NEP activity and modulate Aβ_42_ levels in the brain^11,12^. In the present study, we address how NEP activity is regulated in the signaling cascade downstream of SST.

Using *in vitro* proteomics, we identified ENSA, an endogenous ligand of the ATP-sensitive potassium (K_ATP_) channel ^13,14^, as a negative regulator of NEP downstream of SST signaling. To analyze the role of ENSA *in vivo,* we generated ENSA-deficient mice using genome-editing tools. ENSA deficiency induced NEP activation and reduced Aβ level in the brains of wild-type (WT) and *App* knock-in mice, which develop amyloid plaques without overexpression of *APP* gene containing pathogenic mutations^15^. We also determined that ENSA is a substrate for NEP both *in vitro* and *in vivo*. Consistently, the expression of ENSA was higher in *App* knock-in mice and AD patients than in controls. We also show that diazoxide (Dz), a K_ATP_ channel agonist used for the treatment of hypoglycemia and other conditions^16,17^, improved amyloid pathology and memory impairment in *App* knock-in mice. Our results establish a molecular link between the K_ATP_ channel and NEP activation and provide new insights into the development of alternative strategies to prevent AD.

## Results

### Identification of ENSA as a regulator of NEP activity *in vitro*

We previously developed a method for measuring NEP activity in primary neurons,^10^ and subsequently developed a co-culture system composed of cortical/hippocampal and basal ganglia neurons in a ratio of 9:1, which contain both SSTR1 and SSTR4.^11^ Treatment of mixed culture of neurons from different brain regions of E16-18 C57BL/6Ncr mice, but not of neurons from the individual brain regions, with SST or TT232, a selective agonists for SSTR1 and SSTR4, respectively, elevated NEP activity (Fig. 1a-c). Co-cultured neurons derived from SSTR1 and SSTR4 double knockout (*Sst_1_/Sst_4_* dKO) mice failed to exhibit the SST-induced NEP upregulation (Extended Data Fig. 1a-c). We next examined if neurons from cortical/hippocampal and/or basal ganglia origin generate a secretory factor that activates NEP in the mixed culture neurons. Cultured cortical/hippocampal and basal ganglia neurons were treated separately with SST, and the collected conditioned media added to co-cultured neurons (Fig. 1d). We collected media from SST-treated WT cortical/hippocampal neurons (Media A) and from basal ganglia neurons (Media B), and found that only Media A significantly elevated the NEP activity of co-cultured neurons (Fig. 1e-f). Media A also elevated NEP activity in co-cultured neurons derived from *Sst_1_/Sst_4_* dKO mice (Extended Data Fig. 1d,e), indicating that the NEP-stimulating element in question is secreted by cortical/hippocampal neurons and not basal ganglia neurons, and that NEP activity is upregulated in a manner independent of the SST-SSTR pathway.

**Fig.1.**
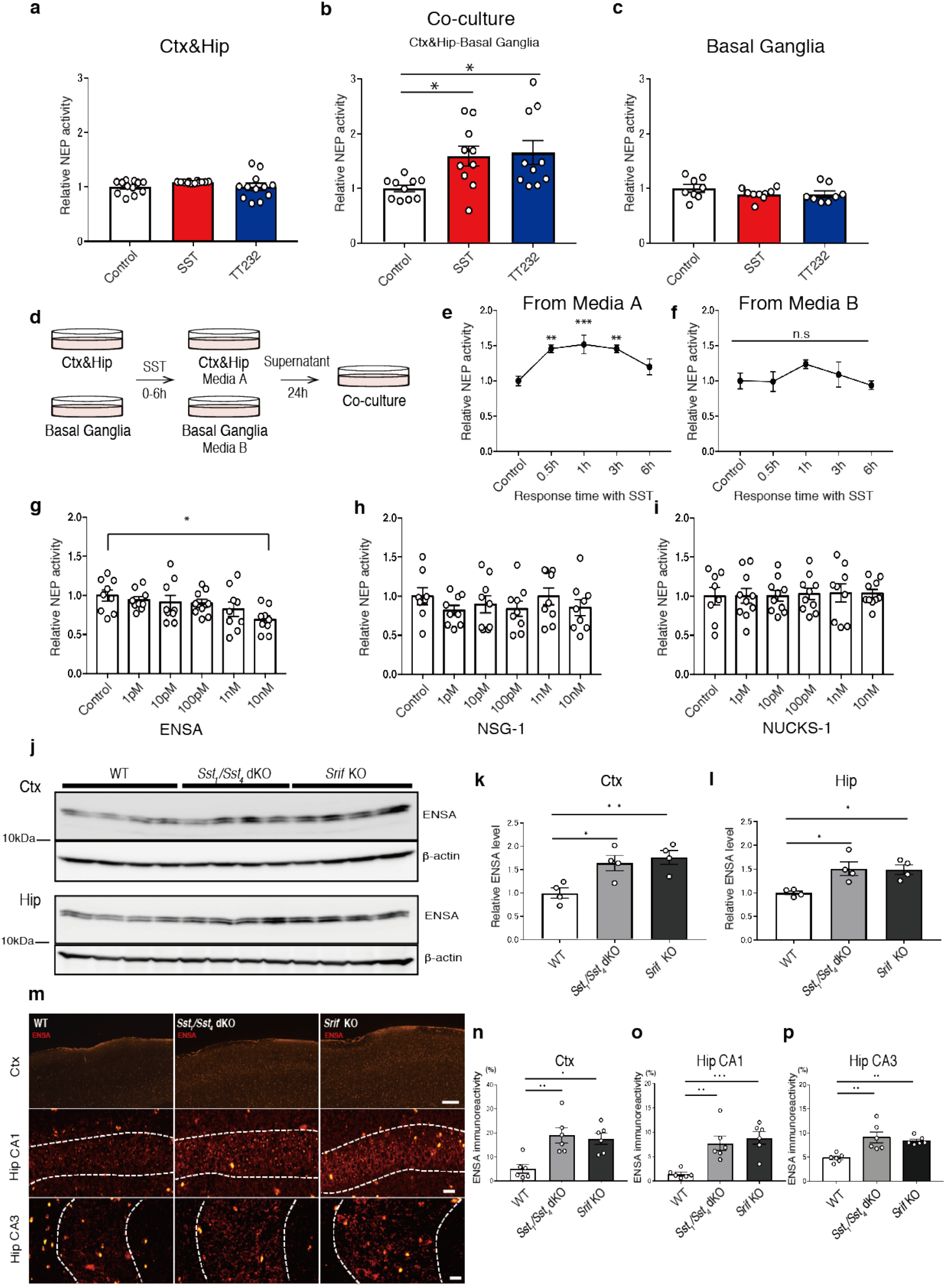
Identification of ENSA as a regulator of NEP activity *in vitro*. **a-c,** NEP activity after treatment of co-cultured cells with 1 µM somatostatin or TT232 for 24 hours. Cortical/hippocampal (Ctx&Hip) neurons (n = 12 wells per treatment), co-cultured neurons (n = 10 wells per treatment), and basal ganglia neurons (n = 8 or 9 wells per treatment) were used. **d-f,** NEP activity in co-cultured neurons after the replacement of the culture medium with conditioned media from Ctx&Hip and basal ganglia neurons treated with 1 µM somatostatin for 0-6 hours. n = 6-10 wells per treatment in co-cultured neurons. **g-i,** NEP activity in co-cultured neurons after incubation with ENSA, NSG-1 and NUCKS-1 recombinant proteins for 24 hours. n = 8-10 wells per treatment in co-cultured neurons. **j-l,** Immunoblotting of ENSA in 3-month-old WT, *Sst_1_/Sst_4_* dKO and *Srif* KO mice (n = 4 for each group). Values indicated in the graph show ENSA band intensities normalized to that of β-actin. **m-p,** Immunostaining of ENSA in the cortices and hippocampal CA1 and CA3 regions from 3-month-old WT, *Sst_1_/Sst_4_* dKO, and *Srif* KO mice (n = 6 for each group). Data represent the mean ±SEM. *\*P*<0.05, *\*\*P*<0.01, *\*\*\*P*<0.001 (one-way ANOVA with Dunnett’s post-hoc test).

A centrifugal filter was used to concentrate NEP activity modulators to the 10-30 kDa molecular weight range (Extended Data Fig. 1f-j), and this fraction was then subjected to LC-MS/MS analysis to identify candidate mediators. Initially, we performed a qualitative comparison between proteins identified in the conditioned media from wild-type primary neurons treated with or without SST and TT232. We also used conditioned media from *Sst_1_/Sst_4_* dKO mice in the LC-MS/MS analysis as a negative control. We then searched for proteins absent or present only in the media of the SST- and TT232-treated WT mixed neurons, but not in the media of *Sst_1_/Sst_4_* dKO neurons. In this way, we identified three candidate proteins: (1) ENSA, (2) neuron-specific protein family member 1 (NSG-1) and (3) nuclear ubiquitous casein and cyclin-dependent substrate 1 (NUCKS-1) (Table 1). To determine which of the candidates is involved in the regulation of NEP activity, we analyzed the effects of corresponding recombinant proteins on NEP activity in co-cultured neurons. Only the recombinant ENSA decreased NEP activity in co-cultured neurons from WT and *Sst_1_/Sst_4_* dKO mice in a dose-dependent manner (Fig. 1g-i and Extended Data Fig. 1k). Consistent with this, treatment of the co-cultured neurons from WT mice, but not from ENSA-deficient mice, with an antibody that specifically neutralizes ENSA (GTX101493) significantly increased NEP activity at a dose of 100 ng/ml (Extended Data Fig. 1l,m).

**Table1.**
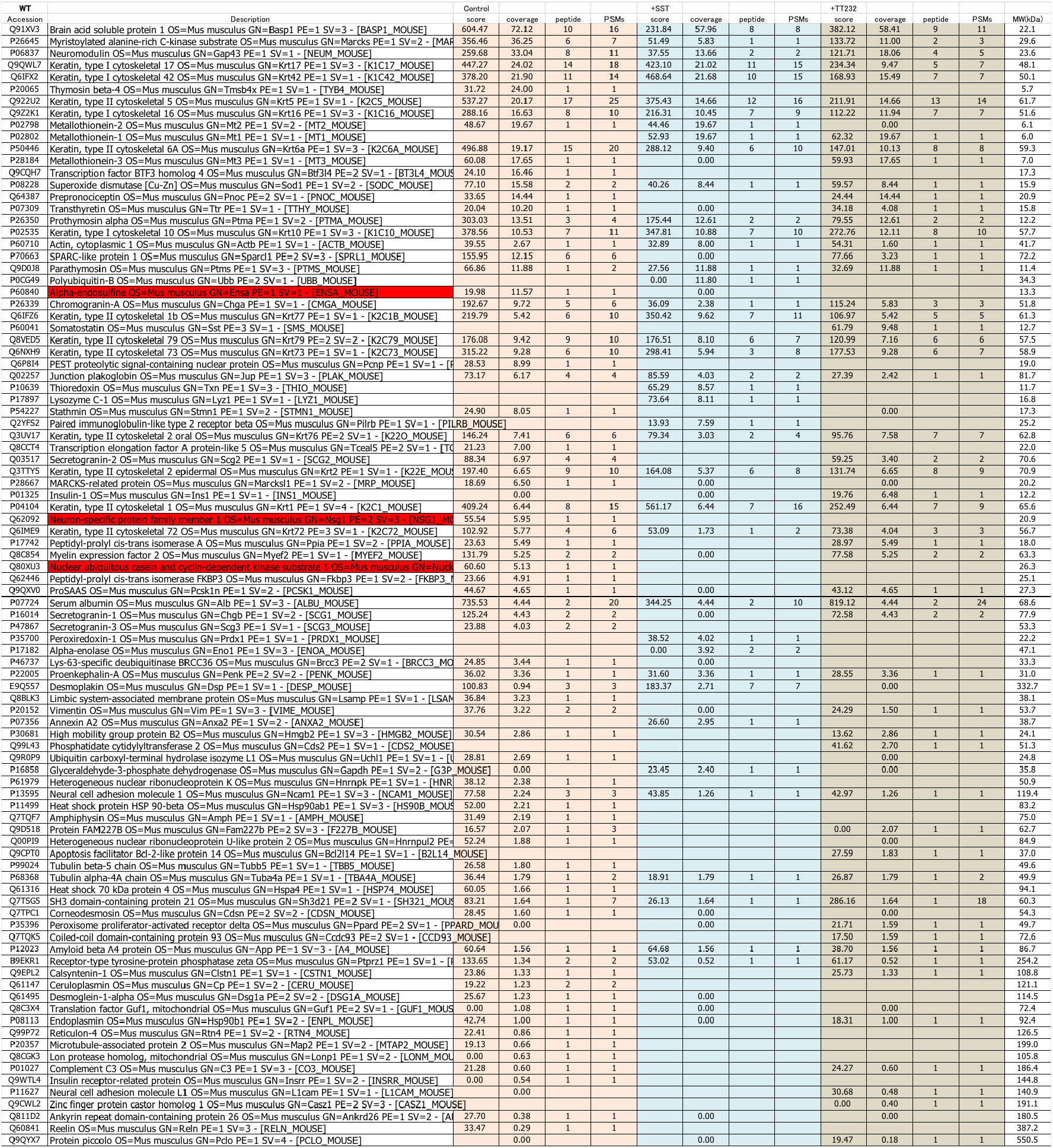
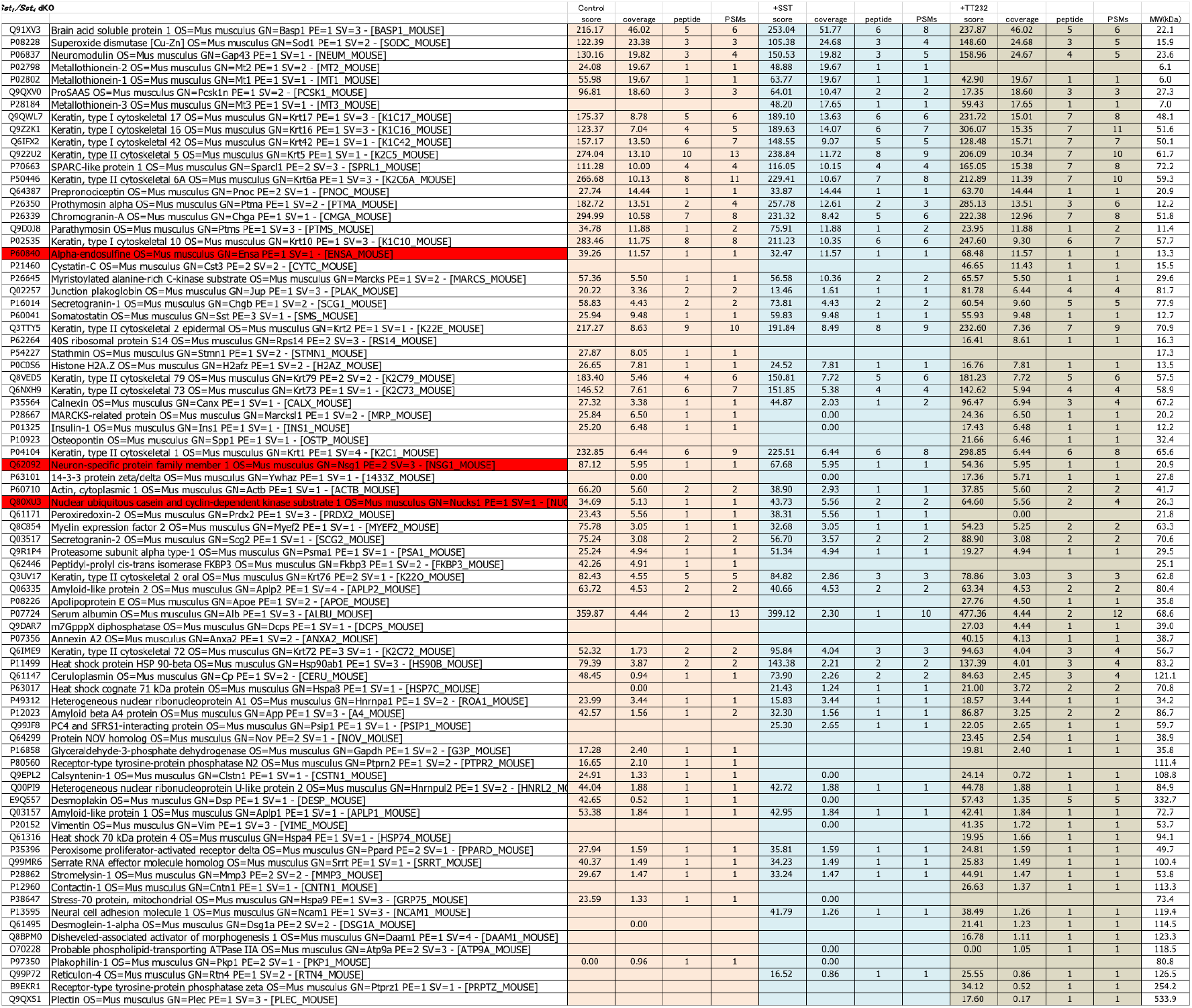
Proteins identified in LC-MS/MS analysis of conditioned media

We next analyzed ENSA levels in the brains of *Sst_1_/Sst_4_* dKO mice and SST knock out (*Srif* KO) mice and found that ENSA levels were significantly elevated in the cortex and hippocampus of both mouse strains (Fig. 1j-l). In addition, immunohistochemical analyses indicated that ENSA-positive signals were heightened in the cortical and hippocampal CA1 and CA3 regions in these animals (Fig. 1m-p). Taken together, these results suggest that ENSA functions both *in vitro* and *in vivo* as a negative NEP regulator downstream of SST signaling.

### Activation of NEP by genetic deficiency of ENSA *in vivo*

ENSA, an endogenous ligand of sulfonylurea receptor 1, regulates the secretion of insulin from pancreatic β cells and is highly expressed in brain and skeletal muscle and at lower levels in the pancreas^13,14^. Although we found that ENSA is a novel negative regulator of NEP *in vitro*, the function of ENSA *in vivo* is largely unknown. We therefore generated ENSA knock out (*Ensa* KO) mice using CRISPR/Cas9 technology. Dual adjacent single-guide RNAs (sgRNAs) were designed that targeted exon 1 of the *Ensa* gene including the initiation codons (Extended Data Fig. 2a). This strategy facilitates CRISPR-mediated genome targeting^18^. We injected sgRNA1-*Ensa*-Exon1 (30 ng/ml) and sgRNA2-*Ensa-*Exon1 (30 ng/ml) together with *Streptococcus pyogenes* Cas9 (SpCas9) mRNA (60 ng/ml) into WT mouse zygotes. Sanger sequencing analysis and PCR-based genotyping indicated the deletion of exon 1, including the initiation codons (Extended Data Fig. 2b,c). Expression of ENSA in homozygous F2 mutant lines, generated by crossbreeding the heterozygous F1 mutant lines, was fully deleted (Extended Data Fig. 2d,e). To assess the off-target effects of CRISPR/Cas9 in the founder, we searched for potential off-target sites using COSMID, with 55 candidate sites being identified (Supplemental Table 1)^19^. Of note, there was no off-target mutation on chromosome 3, which contains the *Ensa* gene. PCR-based genotyping and Sanger sequencing analyses for each candidate site revealed that founder had an off-target mutation in an intergenic region of chromosome 2 which was removed by backcrossing the mutant mice with WT mice (Extended Data Fig. 2f, g).

NEP efficiently degrades Aβ_42_ in the presynaptic region rather than inside secretary vesicles^10^. To determine whether a deficiency of ENSA affects the localization of NEP, we used immunohistochemistry to analyze the expression of NEP and vesicular GABA transporter (VGAT; a presynaptic marker) in the brains of *Ensa* KO mice. We found that NEP signals in the outer molecular layer of the dentate gyrus (OMo) were significantly increased (Fig. 2a,b), and that colocalization of NEP and VGAT these mice was increased in both the lacunosum molecular layer (LM) and OMo (Fig. 2a,c). Next, we measured NEP activity in hippocampal membrane fractions from *Ensa* KO mice and found that a deficiency of ENSA paralleled that of a significantly increased NEP activity (Fig. 2d). We then quantified Aβ_40_ and Aβ_42_ levels in the hippocampi of *Ensa* KO mice by enzyme-linked immunosorbent assay (ELISA) and found that Aβ_42_ levels were significantly reduced compared to those of control mice (Fig. 2f), with Aβ_40_ levels remaining relatively stable (Fig. 2e). This reduction of Aβ_42_ was reproduced in another line of *Ensa* KO mice (*Ensa* KO #2) that was generated by CRISPR/Cas9 with different sgRNAs (Extended Data Fig. 3). These results are in agreement with the effect of somatostatin deficiency^10^. Aβ_42_ levels in ENSA and NEP double knock out (*Ensa/Mme* dKO) mice did not differ from those of single *Mme* KO mice (Fig. 2g), indicating that NEP mediated the reduction of Aβ_42_ in the hippocampi of *Ensa* KO mice.

**Fig.2.**
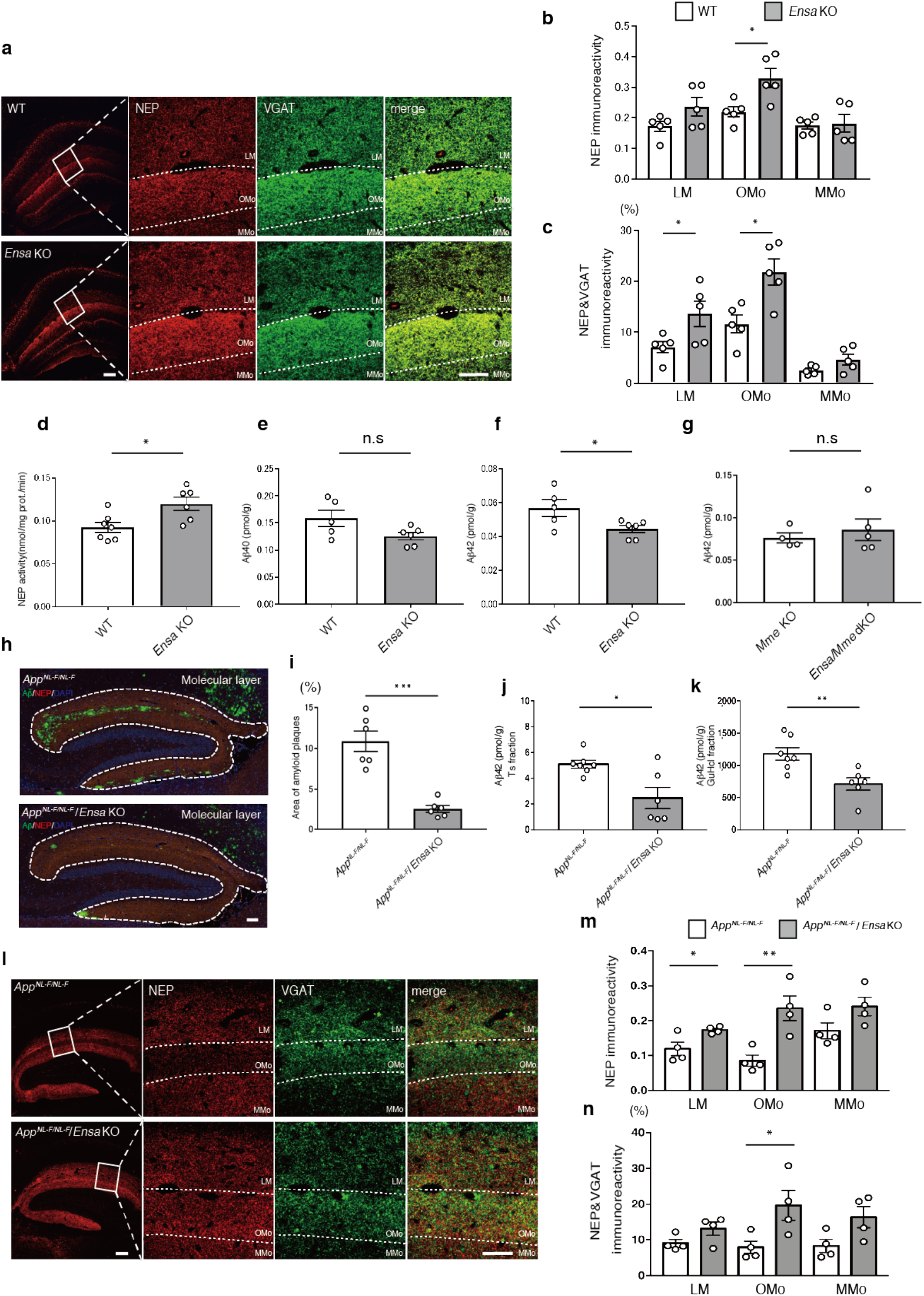
Elevation of NEP activity in *Ensa* KO mice. **a,** Immunostaining of NEP (Red) and VGAT (Green) from hippocampi of 3-month-old WT and *Ensa* KO mice. Scale bar is 100 µm in low magnification image and 50 µm in high-magnification image. **b,** Statistical analysis of NEP immunoreactivity (n = 5 for each group). LM: lacunosum-molecular layer, Omo: Outer molecular layer and MMo: middle molecular layer. **c,** Statistical analysis of co-localized NEP and VGAT signals (n = 5 for each group). **d,** NEP activity in membrane fractions from hippocampi of 3-month-old WT and *Ensa* KO mice (WT: n = 7, *Ensa* KO: n = 6). **e,** Aβ_40_ ELISA of hippocampi of 3-month-old WT and *Ensa* KO mice (WT: n = 5, *Ensa* KO: n = 6). **f,** Aβ_42_ ELISA of hippocampi of 3-month-old WT and *Ensa* KO mice (WT: n = 5, *Ensa* KO: n = 6). **g,** Aβ_42_ ELISA of hippocampi of 3-month-old *Mme* KO mice and *Ensa*/*Mme* dKO (*Mme* KO: n = 4, *Ensa*/*Mme* dKO: n = 5). **h,i,** Immunostaining of Aβ (Green), NEP (Red) and DAPI (blue) from 18-month-old *App^NL-F^* and *App^NL-F^*/*Ensa* KO mice. Statistical analysis of amyloid plaque area in 18-month-old *App^NL-F^* and *App^NL-F^*/*Ensa* KO mice (n = 6 for each group). Scale bar is 100 µm **j,** Aβ_42_ ELISA of Tris-HCl-buffered saline-soluble (Ts) hippocampal fractions from 18-month-old *App^NL-F^* and *App^NL-F^*/*Ensa* KO mice (*App^NL-F^*: n = 7, *App^NL-F^*/*Ensa* KO: n = 6). **k,** Aβ_42_ ELISA of guanidine-HCl-soluble (GuHCl) hippocampal fractions from *App^NL-F^* and *App^NL-F^*/*Ensa* KO mice (*App^NL-F^*: n = 7, *App^NL-F^*/*Ensa* KO: n = 6). **l,** Immunostaining of NEP (Red) and VGAT (Green) in hippocampi of 18-month-old *App^NL-F^* and *App^NL-F^*/*Ensa* KO mice. Scale bar is 100 µm in low-magnification image and 50 µm in high-magnification image. **m,** Statistical analysis of NEP immunoreactivity (n = 4 for each group). **n,** Statistical analysis of co-localized NEP and VGAT signals (n = 4 for each group). Scale bar is 100 µm in low-magnification image and 50 µm in high-magnification image. Data represent the mean ±SEM. *\*P*<0.05, *\*\*P*<0.01, *\*\*\*\*P*<0.0001 (Student’s or Welch’s *t*-test).

We next investigated whether the deficiency of ENSA affected the processing of Aβ production or expression of other Aβ-degrading enzymes. We performed Western blot analysis of full-length APP, its C-terminal fragments generated by α-secretase (CTF-α) and β-secretase (CTF-β), insulin-degrading enzyme (IDE), and endothelin converting enzyme 1 (ECE-1). No significant differences were observed in the expression levels of these proteins and fragments (Extended Data Fig. 2h). Mitogen-activated Protein Kinase/Extracellular Signal-regulated Kinase (ERK1/2) and protein phosphatase 1 (PP1) regulates NEP’s cell surface localization thorough modulation of phosphorylation status in intracellular domain of NEP^20^. The phosphorylation statuses of ERK1/2 at the threonine 202 and tyrosine 204 and PP1 at the threonine 320 residues, which indicate their activity condition respectively^21,22^, however, remained unchanged in *Ensa* KO mice (Extended Data Fig. 4a-c).

To examine the effect of ENSA deficiency on Aβ pathology, we next crossbred *Ensa* KO mice with *App^NL-F/NL-F^* Knock-in (*App^NL-F^*) mice. *App^NL-F^* mice harbor two familial AD-causing mutations (Swedish (KM670/671NL) and Beyreuther/Iberian (I716F)) in the endogenous *App* gene as well as humanized Aβ sequences, and develop amyloid pathology in the hippocampus and cortex from around 6 months of age^15^. The percentage of amyloid plaque deposition in hippocampal molecular layer area was significantly reduced in *App^NL-F^*/*Ensa* KO mice, where NEP expression was elevated (Fig. 2h,i). This result was confirmed in Aβ ELISA experiments on the hippocampi of these mice which showed that Aβ_42_ was significantly decreased (Fig. 2j,k). We consistently found that NEP expression in the LM and OMo of *App^NL-F^*/*Ensa* KO mice was upregulated, particularly in the presynaptic region of OMo (Fig. 2l-n). Taken together, these observations suggest that ENSA is a negative regulator of NEP activity *in vivo* and that a deficiency of ENSA attenuates Aβ pathology by allowing NEP activity to be upregulated.

NEP activity was also measured in cardiac fractions from *Ensa* KO mice given that LCZ696, a dual-acting angiotensin-receptor-neprilysin inhibitor drug, has been approved and is being used to treat heart failure.^23,24^ NEP activity was unaltered in the heart and kidney of *Ensa* KO mice compared to WT controls (Extended Data Fig. 5a,b). Indeed, NEP expression in the kidney was much higher than that in the heart, which is consistent with a previous report stating that the kidney expresses the highest level of NEP among all the mammalian organs^25^.

### Feedback mechanism regulating NEP activity

Previously, several substrates for NEP such as enkephalin, neuropeptide Y and Aβ were identified^26–29^. SST is an endogenous upregulator of NEP and is also degraded by NEP in a substrate-dependent feedback manner^10^. We hypothesized that NEP might also directly degrade ENSA in a similar feedback manner. Co-incubation of recombinant ENSA with NEP resulted in a remarkable decrease in ENSA levels (Fig. 3a). Several NEP inhibitors such as thiorphan, phosphoramidon and EDTA attenuated this effect, indicating that NEP degrades ENSA *in vitro* (Fig. 3a). In contrast, NSG-1 and NUCKS-1 were not cleaved by NEP (Extended Data Fig. 6a). To identify the NEP-mediated cleavage sites in ENSA, we performed MALDI-TOF analysis after incubation of recombinant ENSA with NEP. Several ENSA fragments were detected in the NEP-treated sample, but not in a sample treated in the presence of thiorphan (Fig. 3b, c and Extended Data Fig. 7). We determined the amino acid sequences of these fragments by LC-MS/MS analysis (Supplemental Table 4), and found that NEP partially cleaved ENSA on the amino-terminal side of hydrophobic amino acids in a manner similar to that of other NEP substrates, including Aβ (Fig. 3d).

**Fig.3.**
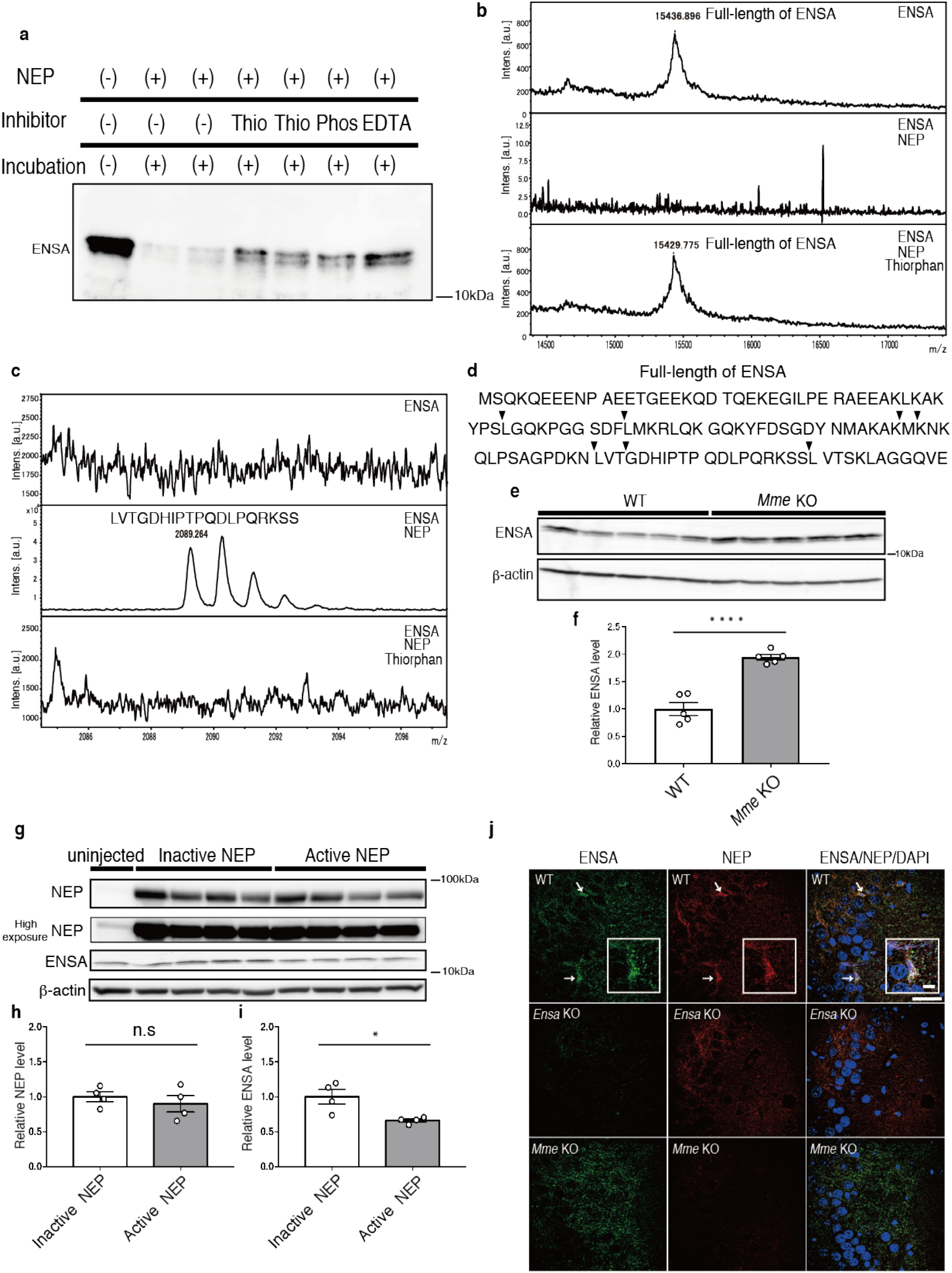
Identification of ENSA as a substrate for NEP. **a,** Immunoblotting of ENSA incubated with or without NEP and mentioned inhibitors for 24 hours at 37°C. Thio: Thiorphan, Phos: Phosphoramidon. **b,** Specific peak of full-length of ENSA after incubation with or without NEP and thiorphan. **c,** Specific peak of cleaved ENSA after incubation with or without NEP and thiorphan. **d,** Sequence of full-length of ENSA. Arrowheads indicate cleavage site by NEP. **e,f,** Immunoblotting of ENSA from hippocampi of 6-month-old WT and *Mme* KO mice. Values indicated in the graph show ENSA band intensities normalized to that of β-actin (n = 5 for each group). **g-i,** Immunoblotting of NEP and ENSA from hippocampi of 3-month-old WT mice after overexpression of active or inactive mutant NEP by SFV gene expression system. Values indicated in the graph show NEP and ENSA band intensities normalized to that of β-actin (n = 4 for each group). **j,** Immunostaining of ENSA (Green), NEP (Red) and DAPI (Blue) in CA3 from 3-month-old WT, *Ensa* KO and *Mme* KO mice. Scale bar is 50 µm in low-magnification image and 10 µm in high-magnification image. White arrows indicate co-localized signals. Data represent the mean ±SEM. *\*P*<0.05, *\*\*\*\*P*<0.0001 (Student’s or Welch’s *t*-test).

ENSA levels in the brains of *Mme* KO mice were subsequently examined by Western blotting and we observed that ENSA was significantly increased in the hippocampi and cortices of these animals (Fig. 3e,f and Extended Data Fig. 6b,c). We then overexpressed WT and inactivated mutant NEP in the hippocampi of WT mice using the Semliki Forest virus (SFV) gene expression system^30^. Exogenously expressed active NEP, but not the inactive mutant, significantly lowered ENSA as well as Aβ_42_ in hippocampi (Fig. 3g-i Extended Data Fig. 6d). We also performed immunohistochemical analyses of ENSA and NEP and found that these two proteins co-localized in the CA3 region (Fig. 3j). These results suggest that NEP directly contributes to the degradation of ENSA *in vivo* and that NEP activity is regulated by a substrate-dependent feedback mechanism.

### Elevated ENSA levels in an AD mouse model and in AD patients

To explore the involvement of ENSA in the etiology of AD, we analyzed ENSA levels in an AD mouse model and in postmortem brain of patients with AD. Western blot analyses revealed that ENSA expression was significantly increased in the cortices and hippocampi of 12- and 24-month-old *App^NL-F^* mice, but not in those of 2-month-old mice which do not yet exhibit amyloid deposition (Fig. 4a-d and Extended Data Fig. 8a-h). In immunohistochemical analyses, ENSA signals in the cerebral cortices and hippocampal CA1 and CA3 regions of *App^NL-F^* mice were also increased at 24 months (Fig. 4e,f). Consistent with these observations, ENSA levels were markedly increased in the cortices of patients with AD (Fig. 4i-l). In contrast, the mRNA levels of ENSA in *App^NL-F^* mice did not differ from those of WT mice at the age of 12 and 24 months (Fig. 4g,h and Extended Data Fig. 8i,j), implying that the proteostasis of ENSA was perturbed in the brains of AD mice. While WT Aβ_42_ inhibited the NEP-mediated degradation of ENSA in a dose-dependent manner *in vitro* (Fig. 4 m,n and Extended Data Fig. 8k,m), mutated Aβ_42_ with the Arctic mutation, that escapes proteolytic degradation by NEP^31^, failed to exert such an effect (Fig. 4 m,n and Extended Data Fig. 8l,m). Consistent with this, another AD mouse model, *App^NL-G-F/NL-G-F^* Knock-in (*App^NL-G-F^*), that harbors the Arctic mutation in addition to the Swedish and Beyreuther/Iberian mutations and exhibits a more aggressive Aβ pathology than *App^NL-F^* mice, showed ENSA levels in the cortex and hippocampus at 6 months (Fig. 4o-r) that were indistinguishable from those of WT mice. ENSA levels in 24-month-old *App^NL-G-F^* mice with more aggressive inflammation than *App^NL-F^* mice were significantly reduced (Extended Data Fig. 8n-r). These results suggest that the elevation of ENSA levels in AD is due to a competitive inhibition between ENSA and Aβ_42_ of NEP-mediated degradation.

**Fig.4.**
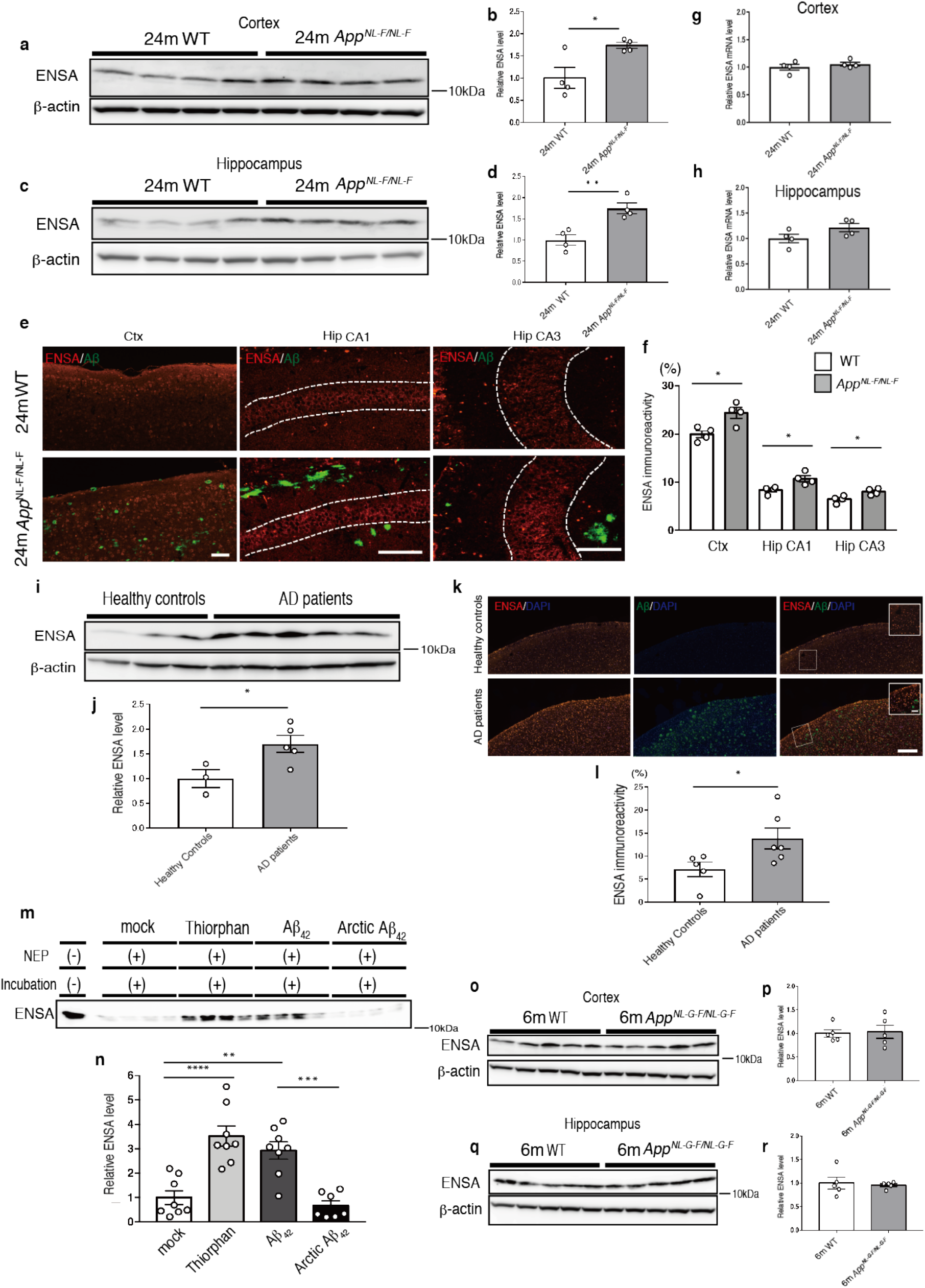
Increased levels of ENSA in AD model mouse and postmortem brain tissue from patients with AD. **a-d,** Immunoblotting of ENSA in cortices and hippocampi of 24-month-old WT and *App^NL-F^* mice. Values indicated in the graph show ENSA band intensities normalized to that of β-actin (n = 4 for each group). **e,f,** Immunostaining of ENSA (Red) and Aβ (Green) in cortex, and hippocampal CA1 and CA3 regions of 24-month-old WT and *App^NL-F^* mice (n = 4 for each group). Scale bar is 100 µm. **g,h,** Semi-quantification of ENSA mRNA levels in cortices and hippocampi of 24-month-old WT and *App^NL-F^* mice. Values indicated in the graph show ENSA band intensities normalized to that of G3PDH (n = 4 for each group). **i,j,** Immunoblotting of ENSA in cortices of healthy controls and AD patients. Values indicated in the graph show ENSA band intensities normalized to that of β-actin (healthy controls: n = 3, AD patients: n = 5). **k,l,** Immunostaining of ENSA in cortices of healthy controls and AD patients (healthy controls: n = 5, AD patients: n = 6). Scale bar is 500 µm in low-magnification image and 100 µm in high-magnification image. **m,n,** Immunostaining of ENSA incubated with or without NEP, Thiorphan, Aβ_42_ and Arctic Aβ_42_ for 24 hours at 37°C (n = 7-8 for each group). **o-r,** Immunoblotting of ENSA in cortices and hippocampi of 6-month-old WT and *App^NL-G-F^* mice. Values indicated in the graph show ENSA band intensities normalized to that of β-actin (n = 4 for each group). In **,b,d,f,j,l** the data represent the mean ±SEM. *\*P*<0.05, *\*\*P*<0.01, (Student’s *t*-test). In **n,** the data represent the mean ±SEM. *\*\*P*<0.01, *\*\*\*\*P*<0.0001 (one-way ANOVA with Turkey’s multiple comparison test). Information concerning human samples is given in Supplemental Table 5.

### Improvement of Aβ pathology and memory function by diazoxide in an AD mouse model

ENSA is known to function as a blocker of the K_ATP_ channel^13^. To investigate whether the K_ATP_ channel modulates NEP activation, we incubated mouse primary neurons with diazoxide (Dz), a K_ATP_ channel agonist, and found that this activated NEP in a dose-dependent manner (Fig. 5a). As Dz has been reported cross the blood brain barrier,^16,17^ we treated WT mice by oral administration of Dz for 1 month. This treatment significantly increased NEP activity in the anterior cortex and hippocampus (Fig. 5b) of these animals, with elevated levels of NEP expression also seen in the anterior cortex (Fig. 5c,d). In line with this, Dz significantly lowered Aβ_42_ levels in the anterior cortex and hippocampus, where NEP was activated (Fig. 5e), whereas Dz treatment had no effect on Aβ_42_ levels in the anterior cortex and hippocampus of *Mme* KO mice (Fig. 5f). These results suggest that Dz decreased Aβ_42_ levels in a NEP-mediated manner.

**Fig.5.**
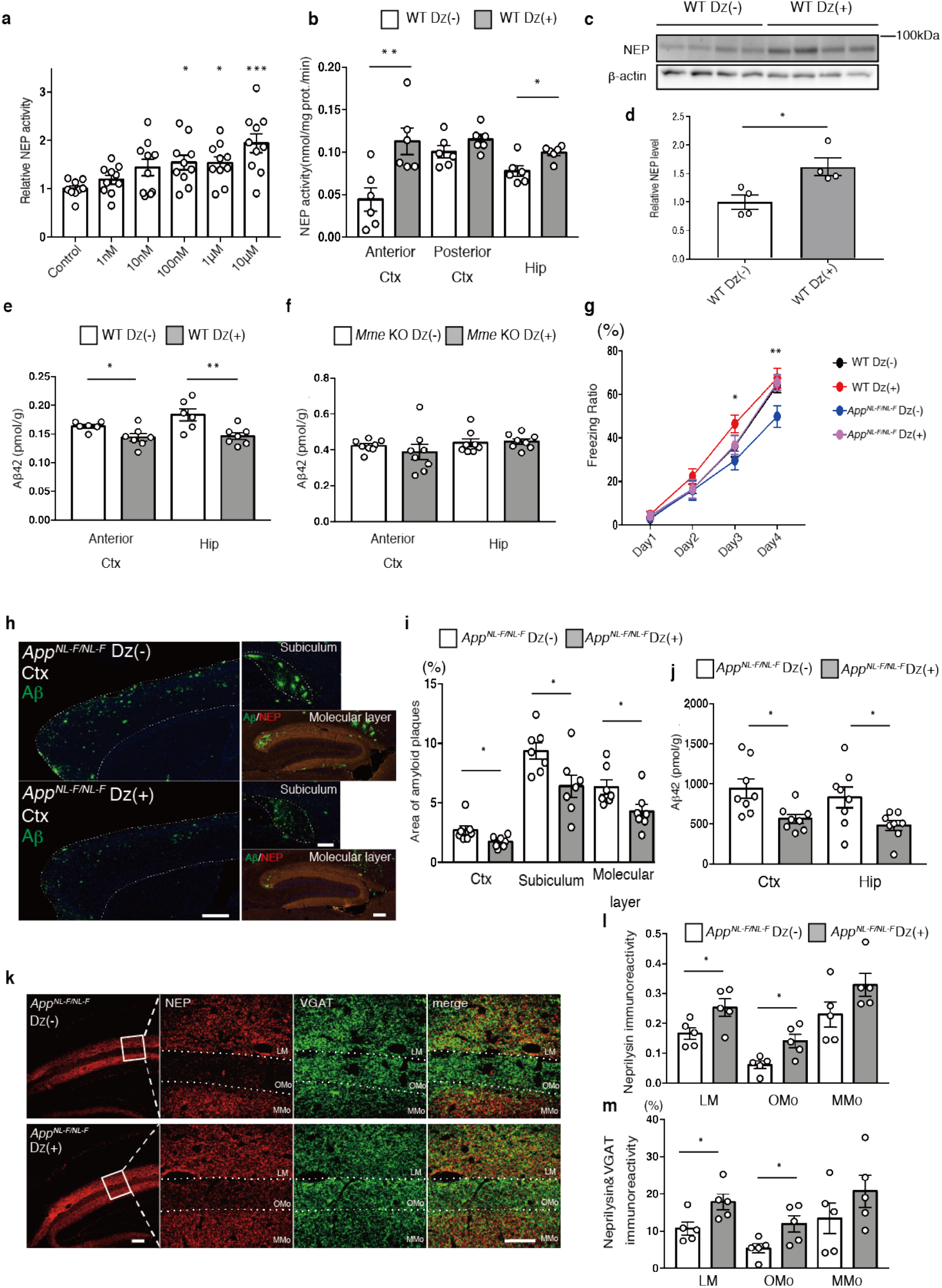
Improvement of Aβ pathology and memory function in *App^NL-F^* mice via enhancement of NEP activity by Dz treatment. **a,** NEP activity after treatment of co-cultured neurons for 24 hours with different doses of diazoxide (Dz) (n = 9-10 for each group). **b,** NEP activity in membrane fractions from anterior cortex (Ctx), posterior Ctx and hippocampus (Hip) of 4-month-old WT mice treated with or without Dz (n = 6 for each group). **c,d,** Immunoblotting of NEP in anterior Ctx of 4-month-old WT mice treated with or without Dz. Values indicated in the graph Values indicated in the graph show NEP band intensities normalized to that of β-actin (n = 4 for each group). **e,** Aβ_42_ ELISA of GuHCl fractions from anterior Ctx and Hip of 4-month-old WT mice with or without Dz (Dz (-): n = 6, Dz (+): n = 7). **f,** Aβ_42_ ELISA of GuHCl fractions from anterior Ctx and Hip of 6-month-old *Mme* KO mice with or without Dz (n=8 for each group). **g,** Freezing ratio of 18-month-old WT and *App^NL-F^* mice treated with or without Dz (WT Dz (-): n = 12, WT Dz (+): n = 13, *App^NL-F^* Dz (-): n = 10, *App^NL-F^* Dz (+): n = 11). **h,i,** Immunostaining of Aβ (Green) and NEP (Red) in Ctx, Subiculum and Molecular layer from 18-month-old *App^NL-F^* with or without Dz (n = 7 for each group). Scale bar in cortical image = 500 µm and hippocampal image = 200 µm. **j,** Aβ_42_ ELISA of GuHCl fractions from cortices and hippocampi of 18-month old *App^NL-F^* with or without Dz (n = 8 for each group). **k,** Immunostaining of NEP (Red) and VGAT (Green) in hippocampi from 18-month old *App^NL-F^* with or without Dz. Scale bar is 100 µm in low-magnification image and 50 µm in high-magnification image. **j,** Statistical analysis of immunoreactivity of NEP (n = 5 for each group). LM: lacunosum-molecular layer, Omo: Outer molecular layer and MMo: middle molecular layer. **k,** Statistical analysis of co-localized signals of NEP and VGAT (n = 5 for each group). In **a,** the data represent the mean ±SEM. *\*P*<0.05, *\*\*\*P*<0.001 (one-way ANOVA with Dunnett’s post-hoc test). In **b, d, e, i, j, l, m,** the data represent the mean ±SEM. *\*P*<0.05, *\*\*P*<0.01, (Student’s *t*-test). In **g,** the data represent the mean ±SEM. On day 3, WT Dz (+) vs *App^NL-F^* Dz (-) *\*P*<0.05. On day 4, WT Dz (-) vs *App^NL-F^* Dz (-) *\*P*<0.05, WT Dz (+) vs *App^NL-F^* Dz (-) *\*\*P*<0.01, *App^NL-F^* Dz (-) vs *App^NL-F^* Dz (+) *\*P*<0.05 (two-way ANOVA with Turkey’s multiple comparison test)

We next investigated the therapeutic effect of Dz on *App^NL-F^* mice by carrying out contextual fear-conditioning tests to assess memory function after 3 months of Dz treatment from the age of 15 months. Dz treatment recovered the freezing ratio of *App^NL-F^* mice to a level comparable to that of WT mice (Fig. 5g). We also performed open field tests to assess the anxiety phenotype as it has been shown that anxiety may affect performance in spatial memory tasks^32^. Dz treatment in WT and *App^NL-F^* mice did not alter the amount of time spent in the central region of the open field maze (Extended Data Fig. 9a), indicating that Dz had no effect on psychological status. In contrast, the Dz treatment improved abnormal spatial memory function in aged *App^NL-F^* mice, and also decreased Aβ plaque deposition in the cortex, subiculum and hippocampal molecular layer (Fig. 5h,i). Aβ_42_ levels in the cortex and hippocampus of these animals were also reduced (Fig. 5j). Immunohistochemical analyses indicated an increase in NEP expression in the anterior cortex of these mice (Extended Data Fig. 9b,c). In addition, co-localized signals of NEP and VGAT were also increased in the presynaptic regions of the hippocampal LM and OMo (Fig. 5k-m). Dz had no effect on behavior (Extended Data Fig. 9d,e) or Aβ pathology in *App^NL-F^*/*Mme* KO mice (Extended Data Fig. 9f,g). Taken together, these results suggest that Dz improves Aβ pathology and memory impairment in *App^NL-F^* mice by upregulating NEP activity.

## Discussion

In the present study, we used *in vitro* and *in vivo* experimental paradigms to identify ENSA as a negative regulator of NEP activity. A genetic deficiency of ENSA increased NEP activity and markedly lowered Aβ_42_ levels. In addition, ENSA was identified as a substrate for NEP, suggesting a potential feedback mechanism for the regulation of NEP activity. Consistently, ENSA levels were found to be increased in AD model mice and AD patients. Moreover, while ENSA functions as an endogenous blocker of the K_ATP_ channel, using Dz as an agonist of the channel prevented Aβ deposition via the activation of NEP and improved memory function in AD model mice. The key findings in this study are schematized in Extended Data Fig.10. While ENSA plays an important role in cell cycle regulation in several cell types^33–35^, its function in the central nervous system however remains largely unknown. Our experiments suggested that ENSA is involved in the Aβ catabolic pathway, achieving its effects via the modulation of NEP activation. A deficiency of ENSA accelerates the translocation of NEP from intracellular secretary vesicles to the presynaptic surface, and while the exact mechanism by which this occurs is unclear, our data indicate that phosphorylation levels of ERK and PP1 in *Ensa* KO mice did not differ from those of WT mice (Extended Data Fig. 4a-c), suggesting that an alternative mechanism regulates the localization of NEP *in vivo*, which remains to be explored.

SST mRNA levels were reported to decrease in brain with aging and in AD^26,36–40^. As such, ENSA, a downstream protein of SST signaling, may be related to the etiology of AD. Indeed, we showed elevation of ENSA levels in the cortices and hippocampi of 12- and 24-moth-old *App^NL-F^* mice as well as in the cortices of AD patients (Fig. 4a-f,i-l and Extended Data Fig. 8e-h). Moreover, *in vitro* and *in vivo* experiments revealed that NEP degraded ENSA as a substrate, suggesting that NEP and ENSA form a negative feedback loop. This hypothesis is based on the fact that opioids and substance P, cell-specific ligands in monocytes and bone marrow cells, respectively^41,42^, regulate NEP via a feedback mechanism. It is possible that Aβ and ENSA compete against each other in the NEP-mediated degradation, additively exacerbating this feedback-loop and inducing a vicious cycle.

A selective agonist of the K_ATP_ channel such as Dz could serve as a beneficial approach to break this vicious cycle given that it is used as a drug for antihypertensive and hypoglycemic properties, and has the potential in the preclinical setting to improve behavioral abnormalities and Aβ pathology in AD^16,17^. A previous study showed that Dz treatment reduced the extracellular accumulation of Aβ in 3xTg mice which display both amyloid and tau pathology due to overexpression of mutated *APP* and *MAPT* genes on a mutant *PSEN1* background^43,44^. The mechanism by which Dz attenuated Aβ plaque deposition was, however, unclear. Our findings indicate that Dz reduced amyloid deposition in *App^NL-F^* mice via the regulation of NEP activity in the anterior cortex and hippocampus. This regional selectivity of NEP regulation by Dz may be dependent on the dopaminergic system in the brain. The K_ATP_ channel is highly expressed in dopaminergic neurons in the midbrain and regulates dopamine release. These neurons project to the frontal cortex and hippocampus^45–50^. Recently we confirmed that dopamine regulates NEP expression and/or activity in the anterior cortex and hippocampus region (N.W, N.K. & T.C.S unpublished data). To further elucidate the mechanism for the regulation of NEP activity, it will be necessary to investigate pathways downstream of ENSA. Likewise, it is important to clarify which K_ATP_ channel subtypes are involved in the regulation of NEP activity in the brain to avoid off-target effects given that different K_ATP_ channel subtypes are expressed in vascular smooth muscle cells, cardiac muscle cells and pancreatic β-cells^51^. In addition to promoting NEP-mediated Aβ degradation, K_ATP_ channel agonists may have beneficial effects in AD. Dz treatment prevents Aβ-induced neurotoxicity induced by oxidative stress and inflammatory damage and also shows neuroprotective effects against apoptosis *in vitro*^52–56^. Compared to Aβ-targeting immunotherapies, synthetic agonists for the K_ATP_ channel are less expensive and would be more acceptable in aging societies around the world. Taken together, we have demonstrated here a new preventive approach at the preclinical stage of AD based on the function of ENSA. This negative regulator of NEP and K_ATP_ channel (via which its effects are mediated) could be a new therapeutic target for lowering Aβ.

## Materials and Methods

### Animals

All animal experiments were conducted according to guidelines of the RIKEN Center for Brain Science. *Srif* KO and *Sst_4_* KO mice were kindly provided by Ute Hochgeschwender, Oklahoma Medical Research Foundation as previously described^11^. *Sst_1_* KO mice were purchased from Jackson laboratory^11^. *Mme* KO mice were used as negative controls^57^. C57BL/6J and ICR mice were used as zygote donors and foster mothers. C57BL/6J mice were also used for backcrossing with *Ensa* KO mice. *App^NL-F^* mice harbor the humanized sequence of Aβ, and the Swedish (KM670/671NL) and Iberian (I716F) mutations, while *App^NL-G-F^* mice harbor the Arctic (E693G) mutation in addition to the humanized sequence of Aβ, and Swedish (KM670/671NL) and Iberian (I716F) mutations as previously described^15^. Male mice were used in all experiments.

### Antibodies

Antibodies used in this research are listed in Supplemental Table 7. The specificity of ENSA antibody was confirmed using the *Ensa* KO mouse.

### Primary neurons

Neurons from the cerebral cortex, hippocampal and basal ganglia regions of brains from embryonic day (E) 16-18 C57BL/6Ncr mice were isolated and cultured. Briefly, brains were excised and placed in culture plates (FALCON) containing neurobasal medium (Thermo Fisher Scientific). The aforementioned brain regions were excised by scalpel and treated with 5 ml of 0.25% trypsin solution (Nacalai tesque 32777-44) at 37°C for 15 minutes. 250 µl of 1% DNase I was added by pipette and mixed. Subsequently, centrifugation was performed at 1500 rpm for 5 minutes and 5 ml of Hank’s buffered salt solution containing 250 µl of 1% DNase I was added to the pellet and incubated in a water bath at 37°C for 5 minutes. An additional 10 ml of Hank’s buffered salt solution was added to the mixture and centrifuged at 1500 rpm for a further 5 minutes. The resulting pellet was added to neurobasal medium with B27 Plus Supplement (Thermo Fisher Scientific 17504044) and 25µM glutamine (Thermo Fisher Scientific 05030-149). The cells were filtered using a cell strainer with 100 µm nylon mesh (Falcon 2360), and seeded on 6- or 96-well plates (Falcon 353046 or Corning 356640). Cortical/hippocampal and basal ganglia neurons were mixed in a 9:1 ratio as co-cultured neurons.

### Neprilysin activity

NEP activity measurements were performed on primary neurons after 15-28 days of in vitro (DIV15-28) culture as previously described^20^. Somatostatin (Peptide institute 4023), TT232 (Tocris 3493), recombinant ENSA (abcam ab92932), recombinant NSG-1 (Creative BioMart NSG1-332H), recombinant NUCKS-1 (Creative BioMart NUCKS1-10956M) and diazoxide (Wako 364-98-7) were added as appropriate concentrations, and cells were incubated for a further 24 hours. Neurons were then incubated with substrate mixture (50 µM suc-Ala-Ala-Phe-MCA (Sigma S8758), 10 nM benzyloxycarbonyl Z-Leu-Leu-Leucinal (Peptide institute 3175-V) and cOmplete EDTA-Free-Protease inhibitor (Roche Diagnostics 4693132) in 0.2 M MES buffer (pH6.5) with or without Thiorphan (Sigma T6031) for 1 hour at 37°C. Following this, 0.1 mM phosphoramidon (Peptide Institute 4082) and 0.1 mg/ml leucine aminopeptidase (Sigma L-5006) were added, and the reaction mixture was incubated at 37°C for a further 30 minutes. 7-Amino- 4-methylcoumarin fluorescence was measured at excitation and emission wavelengths of 380 nm and 460 nm, respectively. Centrifugal 10 and 30 kDa filters (Merck UFC503096, 501096) were used to separate conditioned media obtained from cortical/hippocampal neurons.

### Preparation of membrane fractions from brain tissue

Brain tissues were homogenized in Tris-buffer (50 mM Tris pH 8, 0.25 M sucrose, EDTA-free cOmplete protease inhibitor cocktail (Roche Diagnostics 05056489001)) and centrifuged at 3600 rpm and 4°C for 30 minutes. Collected supernatants were centrifuged at 70,000 rpm at 4°C for 20 minutes. Resultant pellets were solubilized in Tris-buffer containing 1% Triton X-100 and incubated on ice for 1 hour before centrifugation at 70,000 rpm at 4°C for 20 minutes. Protein concentrations of membrane fractions in collected supernatant samples were measured by BCA protein assay kit (Thermo Fisher Scientific 23225).

### LC-MS/MS analysis

50 mM Ammonium bicarbonate, 10% acetonitrile and 20 mM dithiothereitol were added to the conditioned media and incubated for 30 minutes at 56°C. Samples were then treated with 30 mM iodoacetamide and incubated for 30 minutes at 37°C and digested by incubation with 100 ng/µl trypsin overnight at 37°C. Peptide sequences were determined by Q Exactive Orbital Mass Spectrometers (Thermo Fisher Scientific)^58^. We used Proteome Discoverer Software (Thermo Fishier Scientific) for identification of proteins and peptides. Proteins identified in conditioned media are listed in Table 1.

### Preparation for Cas9 and sgRNAs

For synthesis of Cas9 mRNA *in vitro*, plasmid vector pCAG-T3-hCAS-pA (Addgene 48625) was linearized by Sph I, then transcribed with T3 RNA polymerase (Promega) in the presence of Ribo m^7^G Cap Analog (promega) as previously described^59^. The MEGAshortscript T7 (Thermo Fisher Scientific AM1354) and MEGAclear (Thermo Fisher Scientific AM1908) kits were used for *in vitro* transcription of sgRNAs, while the CRISPR Design tool was used for creating sgRNAs^60^. All oligonucleotide sequences used for *in vitro* transcription are listed in Supplemental Table 3.

### Microinjection of mouse zygotes

The SpCas9 mRNA (60 ng/µl) and sgRNAs (30 ng/µl) were injected into the cytoplasm of C57BL/6J zygotes. After incubation at 37℃ for 24 hours, embryos developed to the 2-cell-stage were transplanted into host ICR mice.

### Off-target analysis

Off-target sites that accepted up to three mismatches were determined by COSMID (https://crispr.bme.gatech.edu/)19. Target sites were amplified from tail genomic DNA by PCR using the Ex Taq Polymerase kit (Takara RR001A) with primers listed in Supplemental Table 2. Target sequencing was performed using a DNA sequencer (ABI 3730xl).

### Genotyping

Genomic DNA was extracted from mouse tail using lysis buffer (100 mM Tris pH 8.5, 5 mM EDTA, 0.2% SDS, 200 mM NaCl, 20 µg/µl Proteinase K) and PCR performed using the specific primer set listed in Supplemental Table 6. PCR products were analyzed by MultiNa (Shimadzu) to evaluate the efficiency of the CRISPR-mediated deletion of the *Ensa* gene. Sanger sequencing analyses were conducted using a DNA sequencer (ABI 3730xl).

### Western blot analysis

Mouse brains were homogenized with lysis buffer (50 mM Tris pH 7.6, 0.15 M NaCl and cOmplete protease inhibitor cocktail (Roche Diagnostics 11697498001)) using a Multi-bead shocker MB (Yasui-Kikai). Samples were rotated at 4°C for 1 hour and centrifuged at 15000 rpm for 30 minutes. Supernatants were collected as lysates and then subjected to sodium dodecyl sulfate-polyacrylamide gel electrophoresis (SDS-PAGE) and transferred to a PVDF or nitrocellulose membranes. For detection of ENSA and CTF-APP, membranes were boiled in PBS for 5 minutes, treated with ECL prime blocking buffer (GE healthcare RPN418) for 1 hour and incubated with antibody at 4°C. Dilution ratios of antibodies are listed in Supplemental Table 7. Immunoreactive bands were visualized by ECL Select (GE Healthcare RPN2235) and a LAS-3000 Mini Lumino image analyzer (Fujifilm).

### Co-incubation of ENSA and NEP

25 ng/µl of ENSA were co-incubated with 2.5 ng/µl NEP, 0-500 nM Aβ_42_ (Peptide Institute 4420-s) and arctic Aβ_42_ (Peptide Institute AF-721), 0.1 mM thiorphan, 1 mM phosphoramidon, and 10 mM EDTA at 37°C for 24 hours in 0.2 M MES buffer pH 6.5.

### Immunohistochemical analysis

After deparaffinization of paraffin-embedded mouse brain sections, antigen retrieval was performed by autoclaving at 121°C for 5 minutes. Sections were then treated with 0.3% H_2_O_2_ in methanol to inactivate endogenous peroxidases. Next, sections were rinsed several times with TNT buffer (0.1 M Tris pH 7.5, 0.15 M NaCl, 0.05% Tween20), blocked for 30 min (TSA Biotin System kit), and incubated overnight at 4°C with primary antibody diluted in TNB buffer (0.1 M Tris pH 7.5, 0.15 M NaCl). Sections were rinsed several times and incubated for 1 hour at room temperature with biotinylated secondary antibody (Vector Laboratories). Next, sections were incubated with HRP-conjugated avidin for 30 minutes and tyramide-enhanced FITC or rhodamine for 10 minutes. Finally, sections were treated with DAPI (Cell Signaling Technology 4083S) diluted in TNB buffer before mounting with PermaFluor (Thermo Fisher Scientific TA-030-FM). Sections were scanned on a confocal laser scanning microscope FV-1000 (Olympus) and a NanoZoomer Digital Pathology C9600 (Hamamatsu Photonics) followed by quantification with Metamorph Imaging Software (Molecular Devices) and Definiens Tissue Studio (Definiens).

### Enzyme-linked immunosorbent assay

Mouse cortices were homogenized in TBS buffer (50 mM Tris pH 7.6, 150 mM NaCl, protease inhibitor cocktail) by a Multi-bead shocker (YASUI KIKAI), centrifuged at 70000 rpm for 20 minutes, and supernatants collected as Tris-soluble fractions. Pellets were rinsed with TBS buffer following which 6M guanidine-HCl solution was added and mixed with a Pellet Pestle (KIMBLE). The mixture was then incubated for 1 hour at room temperature. Next, samples were centrifuged at 70000 rpm for 20 minutes and supernatants collected as guanidine-soluble fractions. Tris-soluble fractions and guanidine-soluble fractions were applied to 96-well plates. Aβ_40_ and Aβ_42_ levels were measured with the aid of an Aβ-ELISA kit (Wako 294-62501,294-62601).

### Determination of amino acid sequence of NEP-cleaved ENSA

After co-incubation of ENSA and NEP with or without thiorphan, MALDI-TOF analysis was performed using Autoflex speed (BRUKER) to detect the specific fragment of ENSA cleaved by NEP. LC-MS/MS analysis was then performed to determine the specific amino acid sequences. Data from LC-MS/MS analyses are listed in Supplemental Table 4.

### SFV injection

WT mice (3 months) were used for this experiment. SFV-NEP vectors (active and inactive forms) were developed previously^30^. Mice were anesthetized with a triple mixed anesthetic (Domitor 0.3 mg/Kg, Dormicum 4 mg/kg, Bettlefar 5 mg/kg), with SFV then injected into the bilateral hippocampus (stereotaxic coordinates: anteroposterior, −2.6 mm; mediolateral, ±3.1 mm; dorsoventral, −2.4 mm) in a total volume of 1 μl using a Hamilton syringe (Altair Corporation), at a constant flow rate of 0.1 μl/min using a Legato 130 syringe pump (KD Scientific, Hollistoon, MA). After injection, mice were administrated with Antisedan 3 mg/kg and maintained for 48 hours in cages with free access to food and water.

### RNA extraction and semi-quantitative RT-PCR

Total RNA was extracted from the cortex and hippocampus of brains using RNAiso Plus (Takara 9109) according to the manufacturer’s protocol. Reverse transcription was performed using ReverTra Ace (TOYOBO FSQ-301). Primer pairs are listed in Supplemental Table 6. Semi-quantitative RT-PCR was conducted using a QuantStudio system (Thermo Fisher Scientific).

### Diazoxide treatment of mice

Diazoxide was diluted in drinking water and administered to WT and *App^NL-F^* mice (10 mg/kg/day). For the short-term treatment, diazoxide was administrated to 3-month-old WT mice for 1 month, while in the long-term treatment, diazoxide was administrated to 15-month-old WT and *App^NL-F^* mice for 3 months. After diazoxide treatment for 3 months, mice were subjected to behavioral tests followed by brain dissection.

### Open field test

WT mice and *App^NL-F^* mice were placed individually in a white noise box for at least 1 hour before starting the test. They were then placed in the middle of an open field maze (600×600 mm) and allowed to explore in the area for 10 minutes. The amount of time that mice spent in the central region was measured as an anxiety parameter.

### Contextual fear conditioning test

Before the start of test, the mice were put in the white noise box for at least 1 hour. Subsequently, the mice were placed into a sound-attenuating chamber and allowed to explore the chamber for 5 minutes. The percent freezing time was measured until mice received an electric shock (7.5mA) to the foot after 4 minutes. As a long-term retention test, the same conditioning experiments were repeated daily for 4 days. The training box was cleaned with water and wiped dry with paper toweling before the next mouse was placed in the chamber. Mice were returned to their cages and provided with free access to food and water.

### Human tissues

Brain samples were kindly provided by Dr. John Trojanowski (University of Pennsylvania) in compliance with RIKEN ethics committee guidelines (approval number Wako3 30-4(2)). Other human samples were obtained from Bio Chain and Tissue solutions. All samples are listed in Supplemental Table 5.

### Statistics

All data are shown as the mean ±SEM. For comparisons between two groups, data were analyzed by Student’s or Welch’s *t*-test. For comparisons among more than three groups, we used one-way analysis of variance (ANOVA) followed by Dunnett’s post hoc analysis or Tukey’s post hoc analysis. In the contextual fear conditioning test, we used two-way ANOVA followed by Tukey’s post hoc analysis. All data were analyzed by Prism7 software (San Diego, CA, USA).

## Acknowledgments

We thank Yukiko Nagai, RIKEN Center for Brain Science, for secretarial assistance and the Research Resource Division of RIKEN Center for Brain Science for technical support. Members of the Laboratory for Proteolytic Neuroscience, RIKEN Center for Brain Science, are thanked for helpful discussions. This work was supported in part by research grants from the Strategic Research Program for Brain Sciences and the Japan Agency for Medical Research and Development (AMED) under Grant numbers JP18dm0107070h0003 (T.S) and JP18dm0207001 (T.C.S).

## Author Contributions

N.W., T.S., T.C.S. designed this study. N.W and T.S planned the experiments. N.W., N.K., M.T performed the experiments. N.W analyzed the data and prepared the Figures and Tables. N.W., S.H., H.S. and T.C.S wrote and edited the manuscript. All authors provided feedback and agreed on the final manuscript.

## Competing Interests

The authors declare no competing interests.

## Supplemental data (Extended data)

**Extended data Fig.1.**
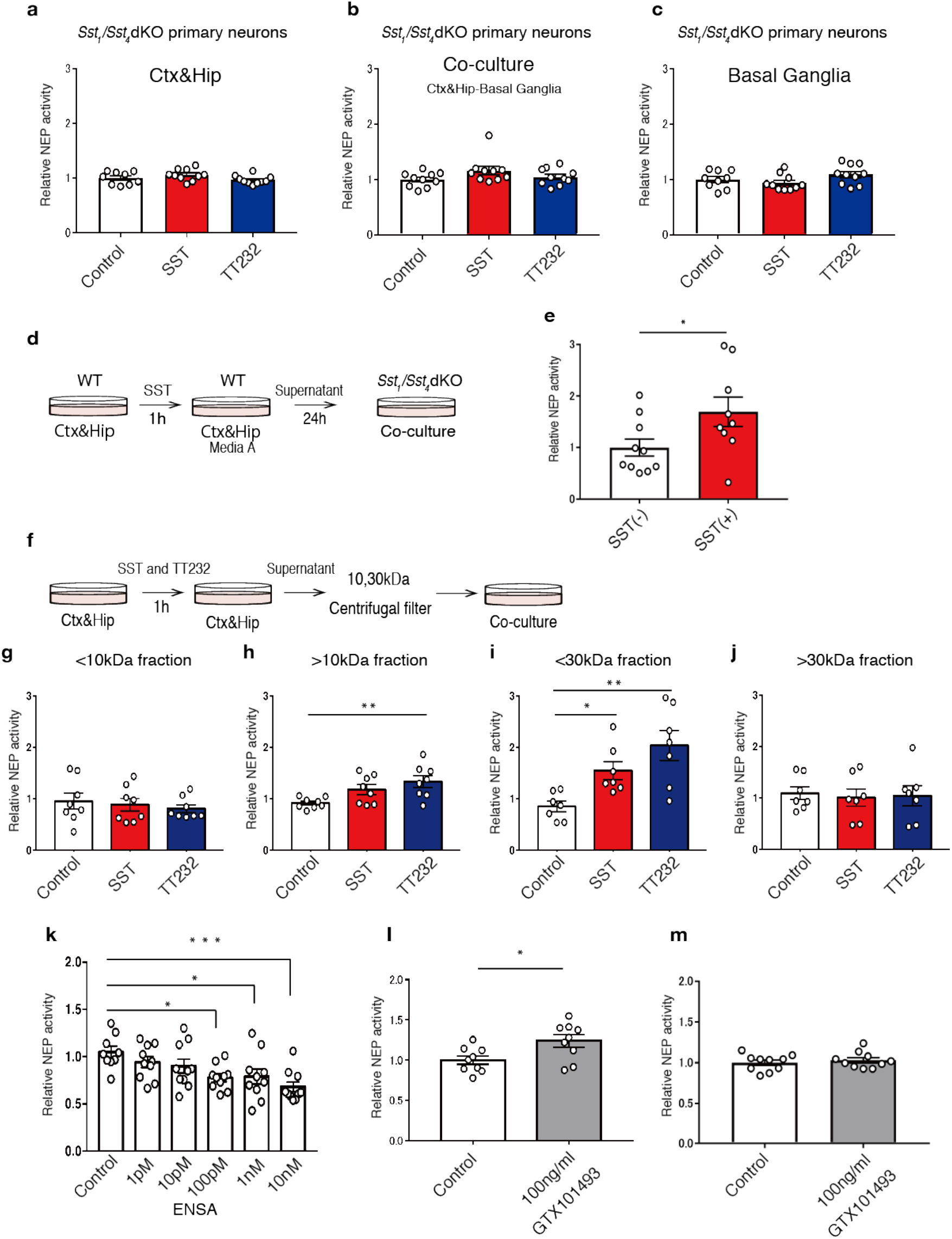
Identification of ENSA as a regulator of NEP activity *in vitro*. **a-c,** NEP activity after treatment of primary neurons derived from *Sst_1_*/*Sst_4_* dKO mice with 1 μM SST or TT232 for 24 hours. Cortical/hippocampal (Ctx&Hip) neurons (n = 9-10 wells per treatment), co-cultured neurons (n = 10 wells per treatment), and basal ganglia neurons (n = 9-10 wells per treatment) were used. **d, e,** NEP activity of co-cultured neurons from *Sst_1_*/*Sst_4_* dKO mice after replacement of the culture medium with conditioned media derived from SST-treated Ctx&Hip neurons from WT mice (n = 9-10 for each group). **f-j,** NEP activity of co-cultured neurons after replacement of the culture medium with separated conditioned media from Ctx&Hip neurons treated with SST or TT232. 10 and 30 kDa centrifugal filters were used for the separation (n = 7-8 for each group). **k,** NEP activity of co-cultured neurons derived from *Sst_1_*/*Sst_4_* dKO mice after treatment with different doses of recombinant ENSA protein (n = 9-10 for each group). **l,** NEP activity of co-cultured neurons after neutralizing ENSA with 100 ng/ml of ENSA-specific antibody (GTX101493) (n = 9 for each group). **g,** NEP activity of co-cultured neurons derived from *Ensa* KO mice after neutralizing ENSA with 100 ng/ml ENSA-specific antibody (GTX101393) (n = 10 for each group). Information concerning *Ensa* KO mice is given in Extended Data Fig.2. In **h, i, k,** the data represent the mean ±SEM. *\*P*<0.05, *\*\*P*<0.01, *\*\*\*P*<0.001 (one-way ANOVA with Dunnett’s post-hoc test). In **e, l,** the data represent the mean ±SEM. *\*P*<0.05 (Student’s *t*-test).

**Extended data Fig.2.**
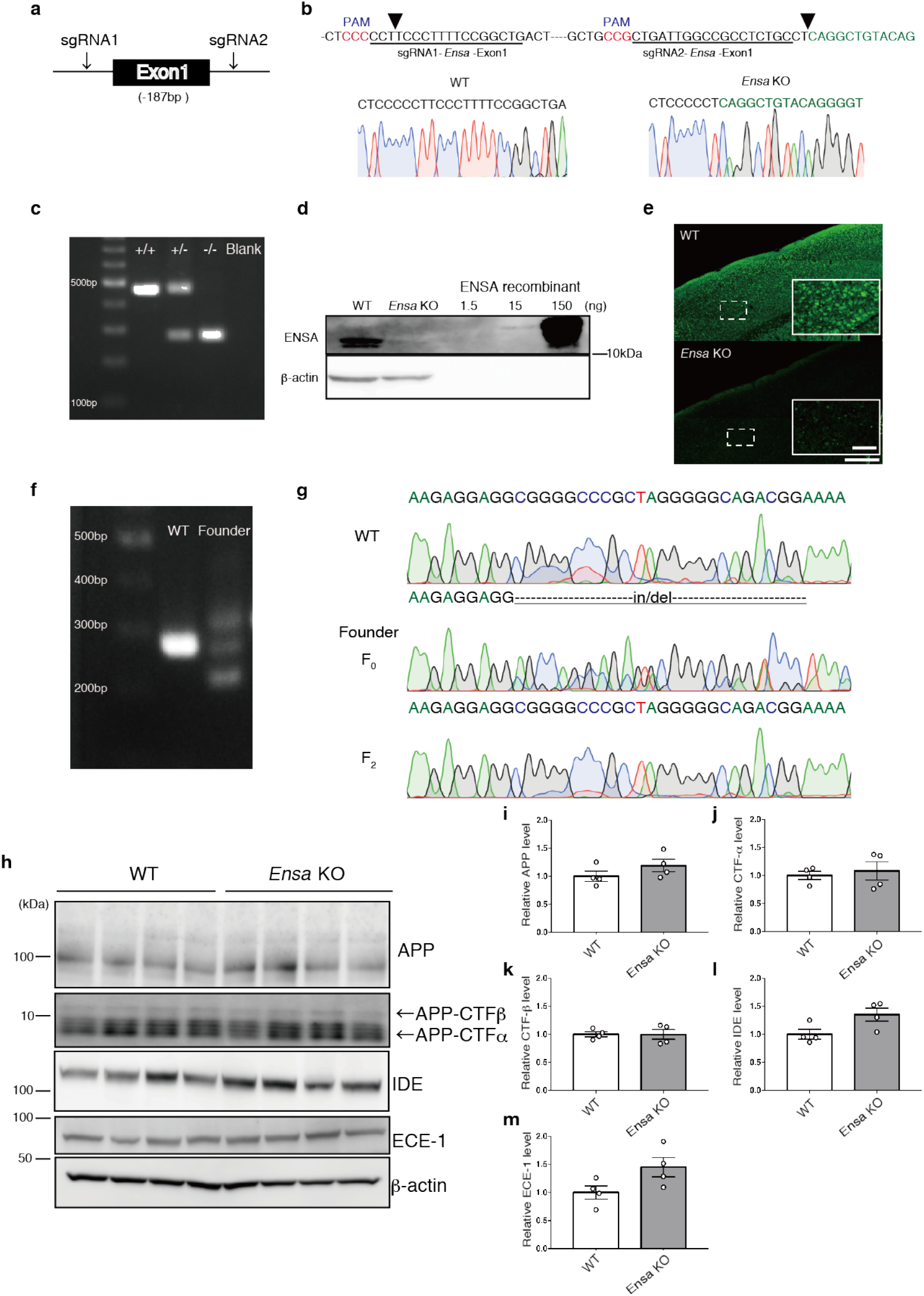
Generation of ENSA-deficient mouse using CRISPR/Cas9. **a,** Schematic image for CRISPR/Cas9-mediated ENSA deficiency. **b,** Sanger sequence chromatograms near exon 1 of *Ensa* gene in WT and *Ensa* KO mice. Arrowheads show Cas9 cleavage sites. **c,** PCR-based genotyping results of WT, heterozygous and homozygous *Ensa* KO mice. Genotyping was performed using mouse tail genome. **d,** Immunoblotting of ENSA in WT and *Ensa* KO mice. **e,** Immunostaining of ENSA in WT and *Ensa* KO mice. ENSA immunoreactivity is absent in *Ensa* KO mice. Scale bar = 500 µm. Inset Scale bar = 50µm. **f,** PCR-based genotyping results of off-target sites in WT and founder *Ensa* KO mice. Genotyping was performed using mouse tail genome. **g,** Sanger sequence chromatograms of off-target sites in WT, founder mouse and F2 *Ensa* KO mouse. **h,** Immunoblotting of APP, CTFs, IDE and ECE-1 in 3-month-old WT and *Ensa* KO mice. **i-m,** Values indicated in graphs show band intensities for APP, CTFs, IDE and ECE-1 normalized to that of β-actin (n = 4 for each group). Results are expressed as the mean ±SEM.

**Extended data Fig.3.**
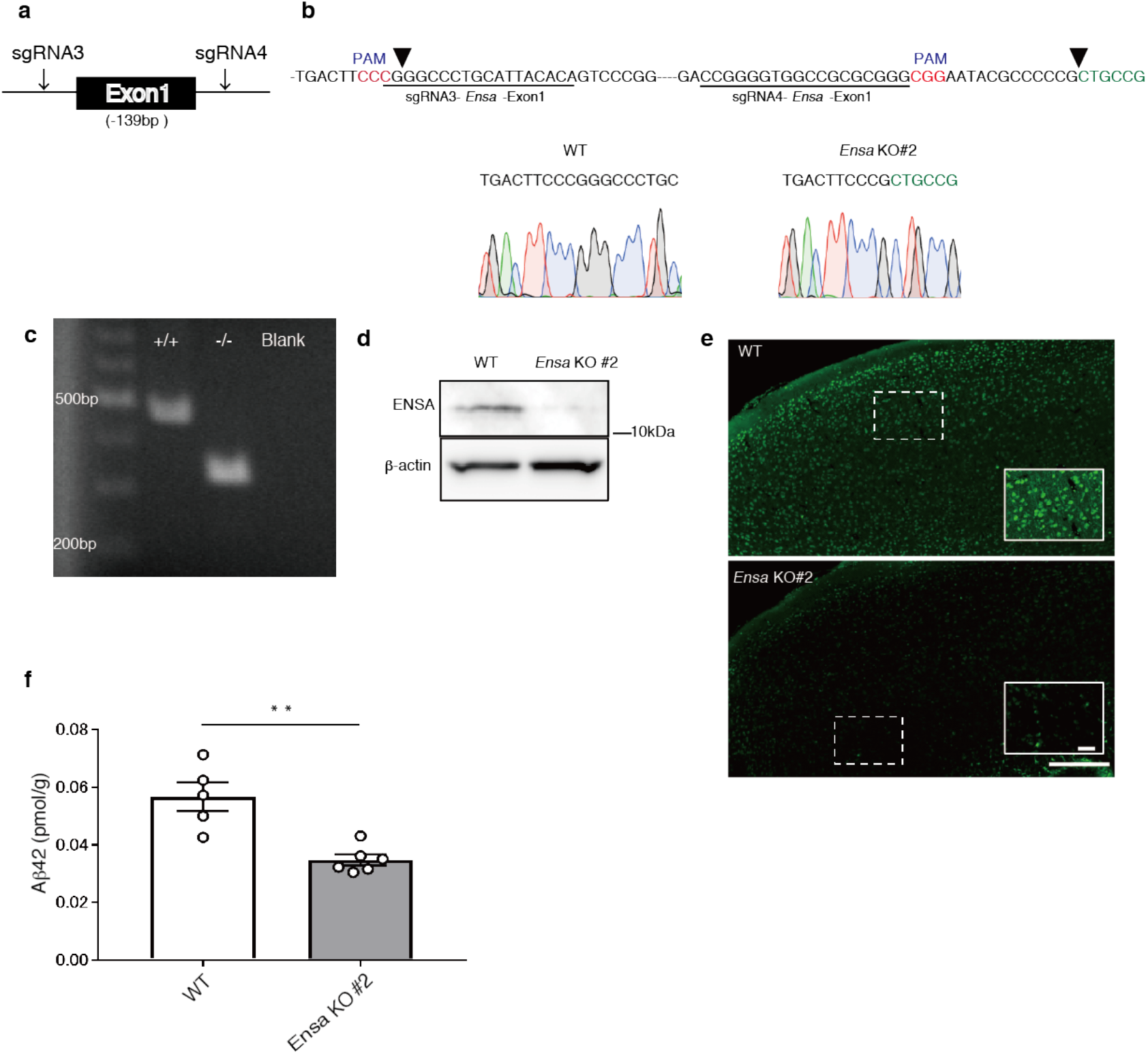
Generation of 2nd line ENSA-deficient mouse using CRISPR/Cas9. **a,** Schematic image for CRISPR/Cas9-mediated ENSA deficiency. **b,** Sanger sequence chromatograms near exon 1 of *Ensa* gene in WT and *Ensa* KO #2 mice. Arrowheads show cleavage sites by Cas9. **c,** PCR-based genotyping results of WT and *Ensa* KO #2 mice. Genotyping was performed using mouse tail genome. **d,** Immunoblotting of ENSA in WT and *Ensa* KO #2 mice. **e,** Immunostaining of ENSA in WT and *Ensa* KO #2 mice. ENSA immunoreactivity was absent in *Ensa* KO #2 mice. Scale bar = 500 µm. Inset scale bar = 50 µm. **f,** Aβ_42_ ELISA of hippocampi from 3-month-old WT and *Ensa* KO #2 mice (WT: n = 5, *Ensa* KO #2: n = 6). Results are expressed as the mean ±SEM. *\*\*P*<0.01 (Student’s *t*-test).

**Extended data Fig.4.**
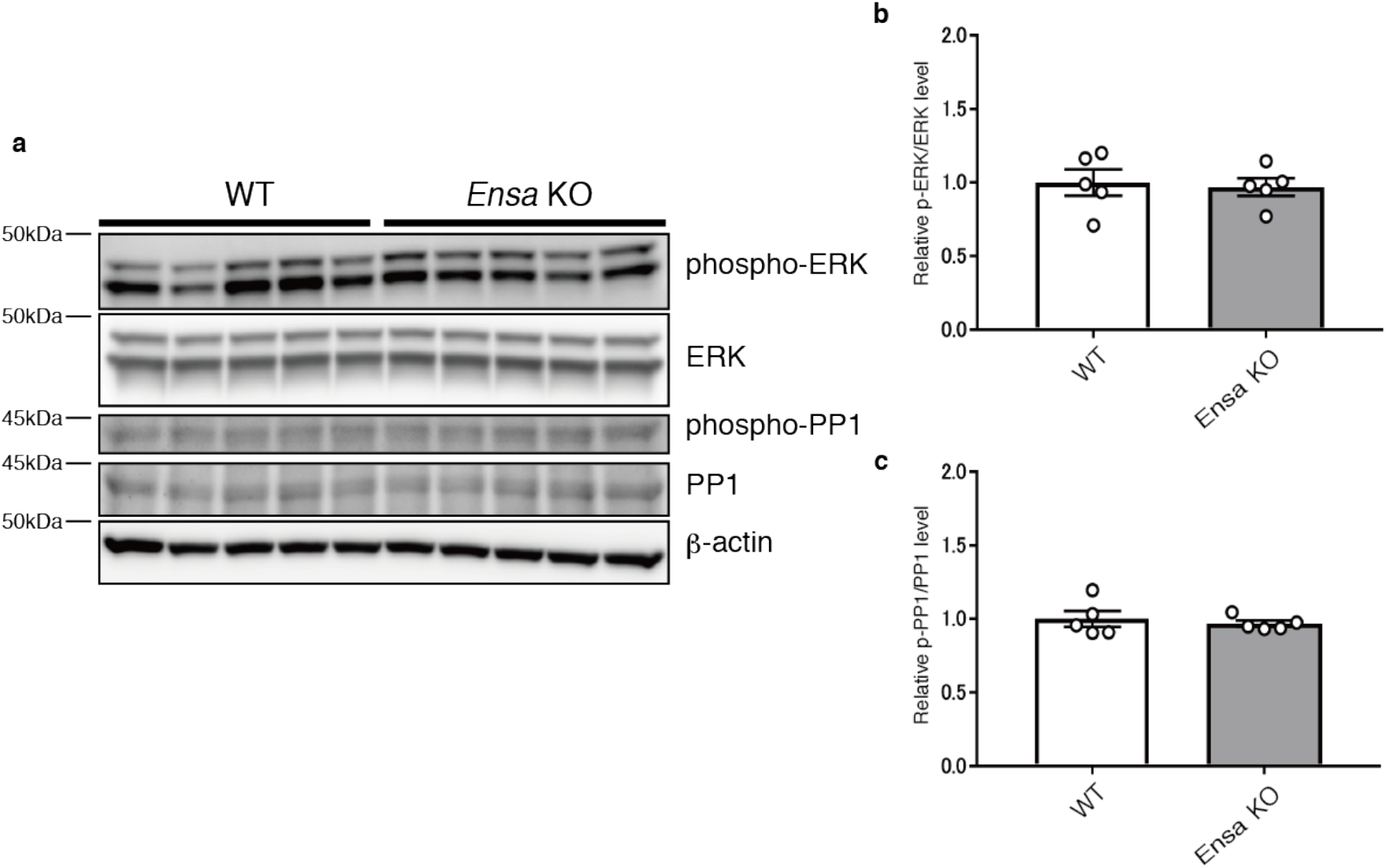
Mechanism for translocalization of NEP in *Ensa* KO mice. **a-c,** Immunoblotting of phospho-ERK, ERK, phospho-PP1 and PP1 in hippocampi of 3-month-old ENSA KO mice. Values indicated in **b** show phospho-ERK band intensities normalized to those of ERK and in **c**, phospho-PP1 band intensity normalized to that of PP1 (n = 5 for each group). Results are expressed as the mean ±SEM.

**Extended data Fig.5.**
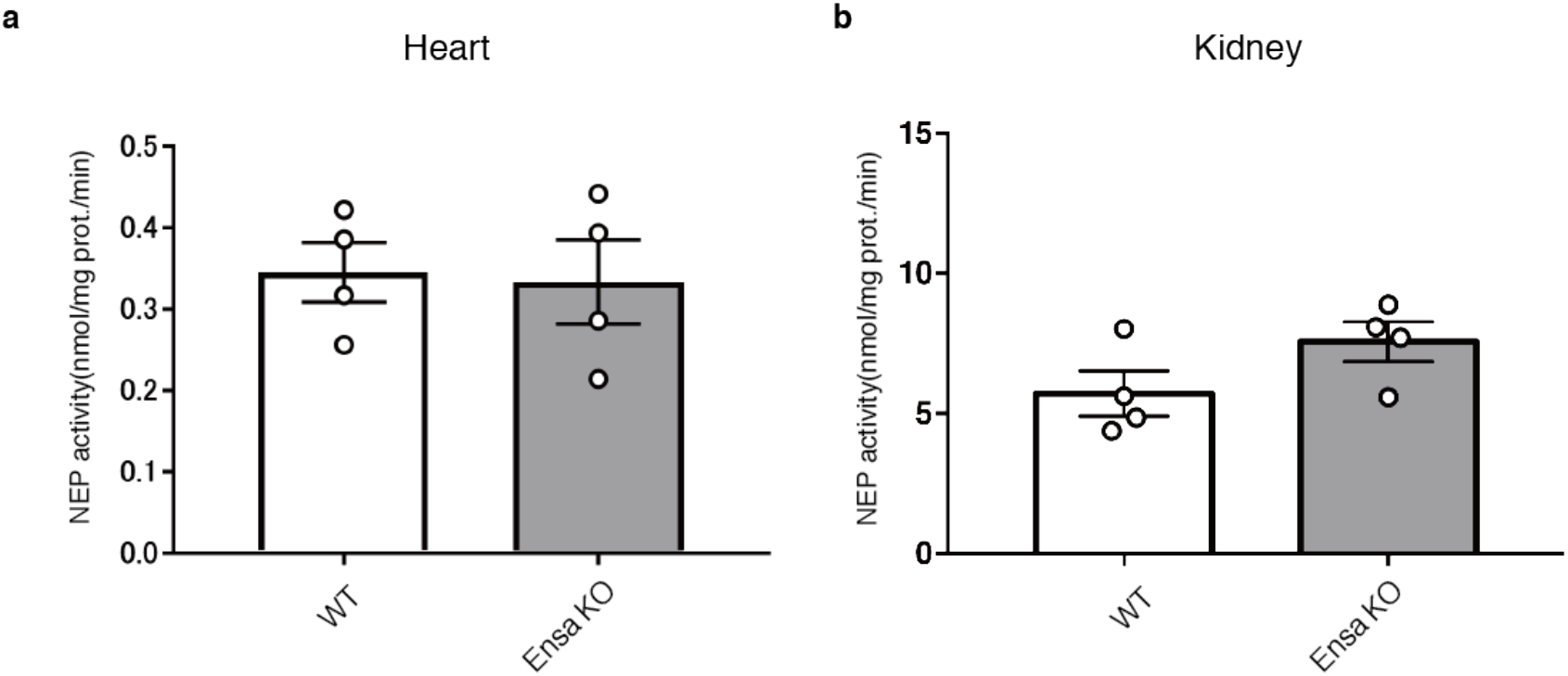
Cardiac and renal NEP activity in *Ensa* KO mice. **a,** NEP activity in membrane fractions from cardiac tissue of 3-month-old WT and *Ensa* KO mice (n = 4 for each group). **b,** NEP activity in the membrane fractions from kidney tissue of 3-month-old WT and *Ensa* KO mice (n = 4 for each group). Results are expressed as the mean ±SEM.

**Extended data Fig.6.**
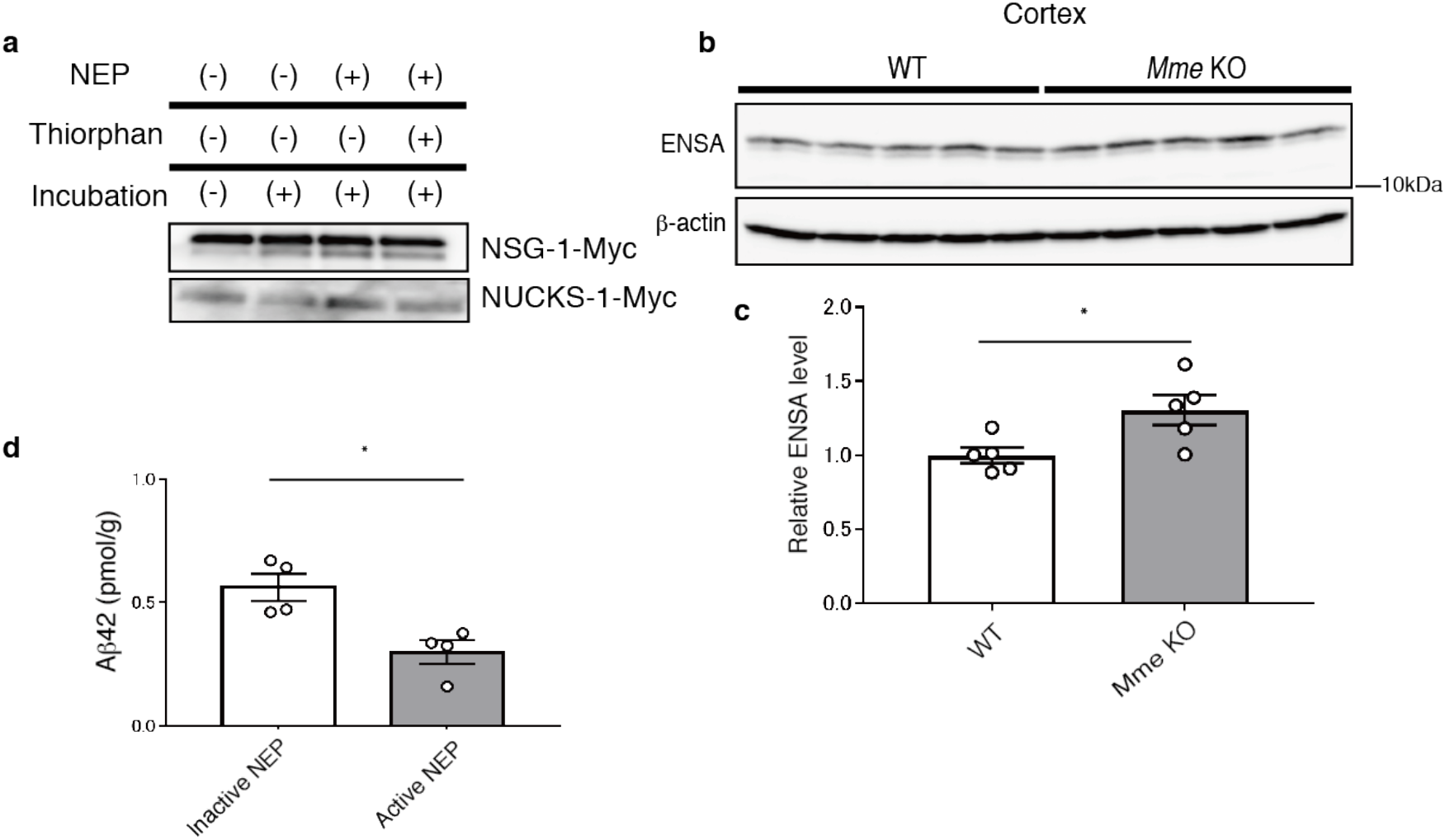
Identification of ENSA as a substrate for NEP. **a,** Immunoblotting of NSG-1 or NUCKS-1 incubated with or without NEP and thiorphan for 24 hours at 37°C. The specific band is detected by Myc-tag antibody. **b,** Immunoblotting of ENSA in cortices from 6-month-old WT and *Mme* KO mice. Values indicated in the graph show the intensity of ENSA bands normalized to that of β-actin (n = 5 for each group). **c,** Aβ_42_ ELISA of Tris-HCl-buffered saline-soluble fractions containing 1% Triton-X from hippocampi of WT mice after overexpression of active or inactive mutant NEP by the SFV gene expression system (n = 4 for each group). Results are expressed as the mean ±SEM. *\*P*<0.05 (Student’s *t*-test).

**Extended data Fig.7.**
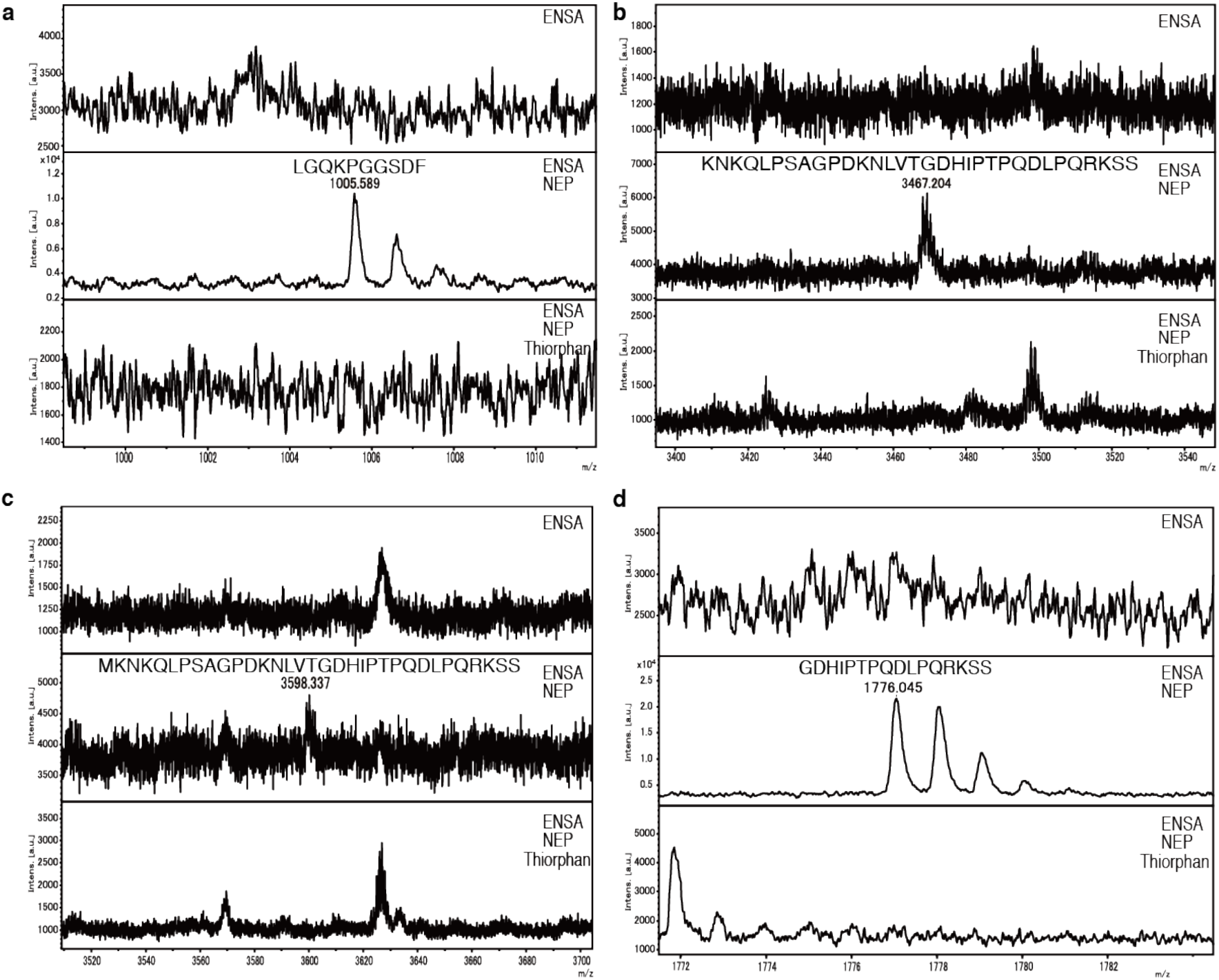
Specific peaks of ENSA cleaved by NEP. **a-d,** MALDI-TOF analyses showing specific peaks of cleaved ENSA after incubation in the presence or absence of NEP and thiorphan for 24 hours at 37°C. LC-MS/MS analysis was used to determine specific amino acid sequences (Supplemental Table 3).

**Extended data Fig.8.**
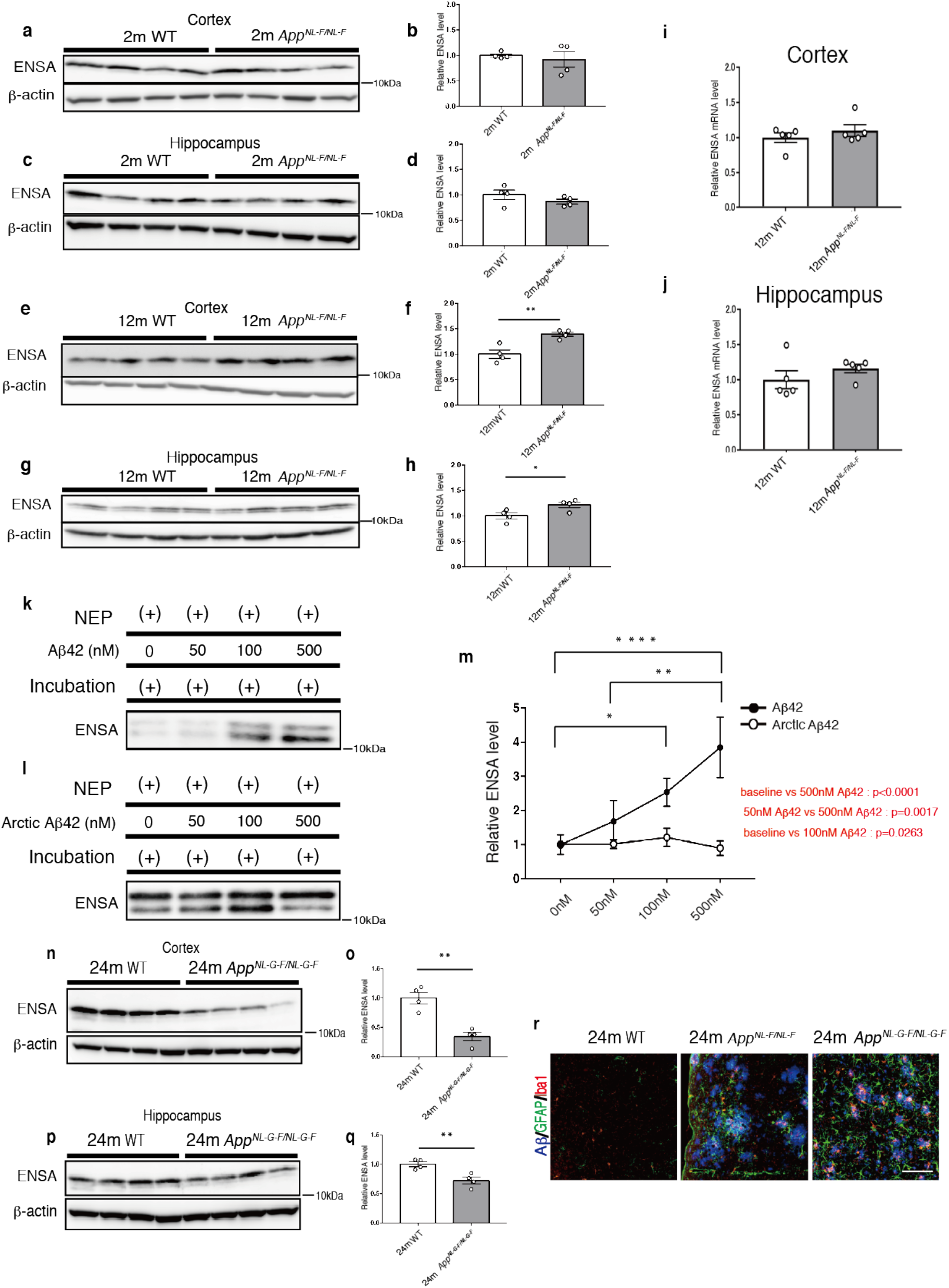
ENSA levels in *App^NL-F^* and *App^NL-G-F^* mice. **a-d,** Immunoblots of ENSA in cortices and hippocampi of 2-month-old WT and *App^NL-F^* mice. Values indicated in the graph show the intensities of ENSA bands normalized to that of β-actin (n = 4 for each group). **e-h,** Immunoblots of ENSA in cortices and hippocampi from 12-month-old WT and *App^NL-F^* mice. Values indicated in the graphs show the intensities of ENSA bands normalized to that of β-actin (n = 4 for each group). **i,j,** Semi-quantification of ENSA mRNA levels in cortices and hippocampi of WT and *App^NL-F^* mice at 12 months. Values indicated in the graphs show ENSA levels normalized to that of G3PDH (n = 4 for each group). **k-m,** Immmunoblots of ENSA incubated with NEP and specified levels of Aβ_42_ or Arctic Aβ_42_ (n = 4 for each group). **n-q,** Immunoblots of ENSA in cortices and hippocampi of 24-month-old WT and *App^NL-G-F^* mice. Values indicated in the graph show ENSA band intensities normalized to that of β-actin (n = 4 for each group). r, Immunostaining of Aβ, GFAP and Iba1 in 24-month-old WT, *App^NL-F^* and *App^NL-G-F^* mice. Scale bar = 100 µm. In **f, h, o, q,** results are expressed as the mean ±SEM. *\*P*<0.05, *\*\*P*<0.01, (Student’s *t*-test). In **m,** results are expressed as the mean ±SEM. *\*P*<0.05, *\*\*P*<0.01, *\*\*\*\*P*<0.0001 (two-way ANOVA with Turkey’s multiple comparison test).

**Extended data Fig.9.**
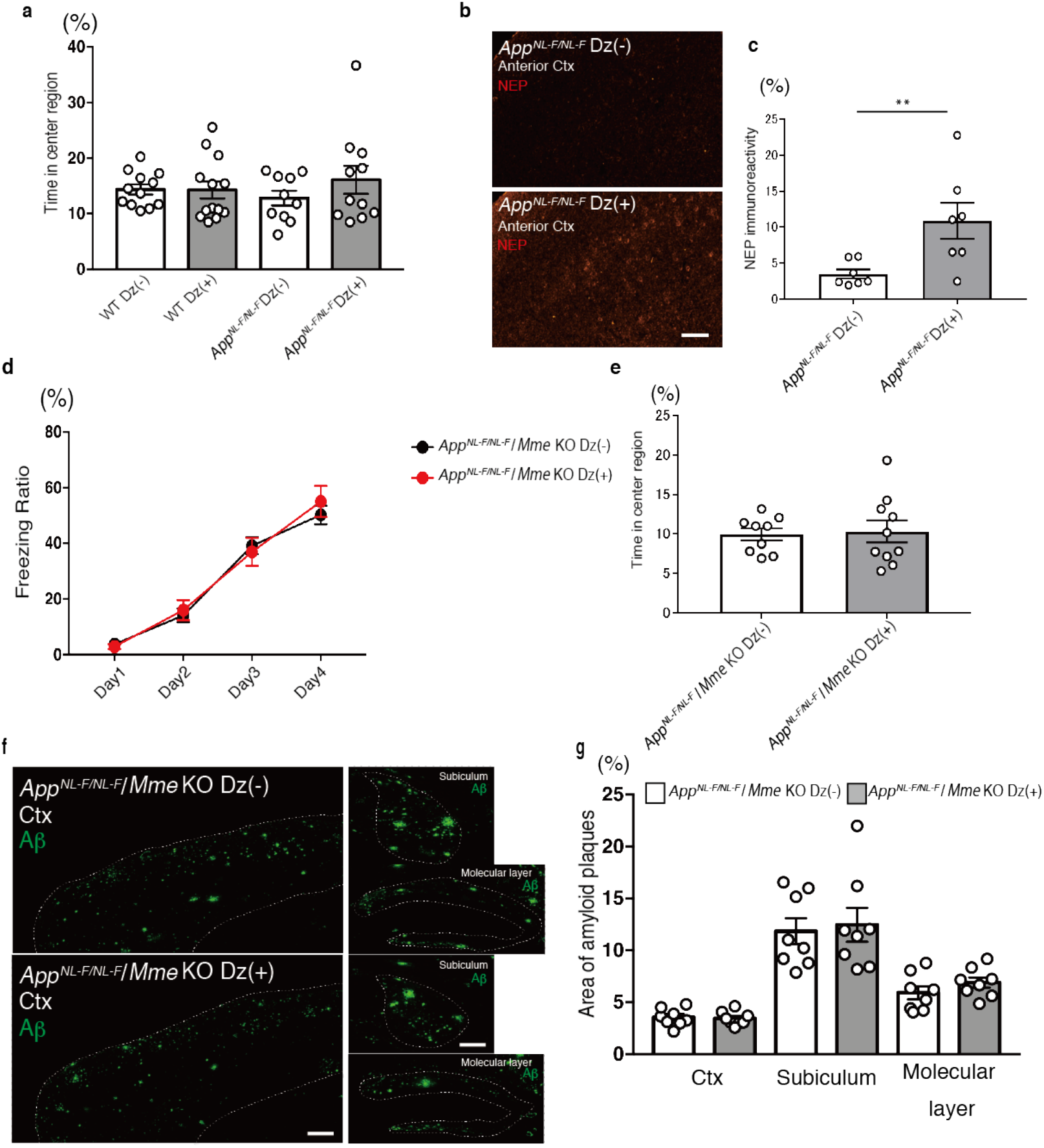
Improvement of Aβ pathology in AD model mouse via enhancement of NEP activity by diazoxide treatment. a, Statistical analysis of open field test to measure time in central region. 18-month-old WT and *App^NL-F^* mice were treated with or without Dz for 3 months (WT Dz (-): n = 12, WT Dz (+): n = 13, *App^NL-F^* Dz (-): n = 10, *App^NL-F^* Dz (+): n = 11). **b,c,** Immunostaining of NEP in cortices of 18-month-old WT and *App^NL-F^* mice treated with or without Dz for 3 months (n = 7 for each group). Scale bar = 500 µm. **d,** Freezing ratio of 15-month-old *App^NL-F^*/*Mme* KO mice treated with or without Dz for 3 months (*App^NL-F^*/*Mme* KO Dz (-): n = 9, *App^NL-F^*/*Mme* KO Dz (+): n = 10). **e,** Statistical analysis of open field test to measure time in central region of maze. 15-month-old *App^NL-F^*/*Mme* KO were treated with or without Dz for 3 months (*App^NL-F^*/*Mme* KO Dz (-): n = 9, *App^NL-F^*/*Mme* KO Dz (+): n = 10). **f,** Immunostaining of Aβ (Green) in cortex, subiculum and molecular layer of 15-month-old *App^NL-F^*/*Mme* KO mice with or without Dz for 3 months. Scale bar in cortical image = 500 µm and in hippocampal image = 200µm. **g,** Statistical analysis of amyloid plaque area in 15-month old *App^NL-F^*/*Mme* KO treated with or without Dz for 3 months (n = 8 for each group). Results expressed as the mean ±SEM. *\*\*P*<0.01 (Mann-Whitney test).

**Extended data Fig.10.**
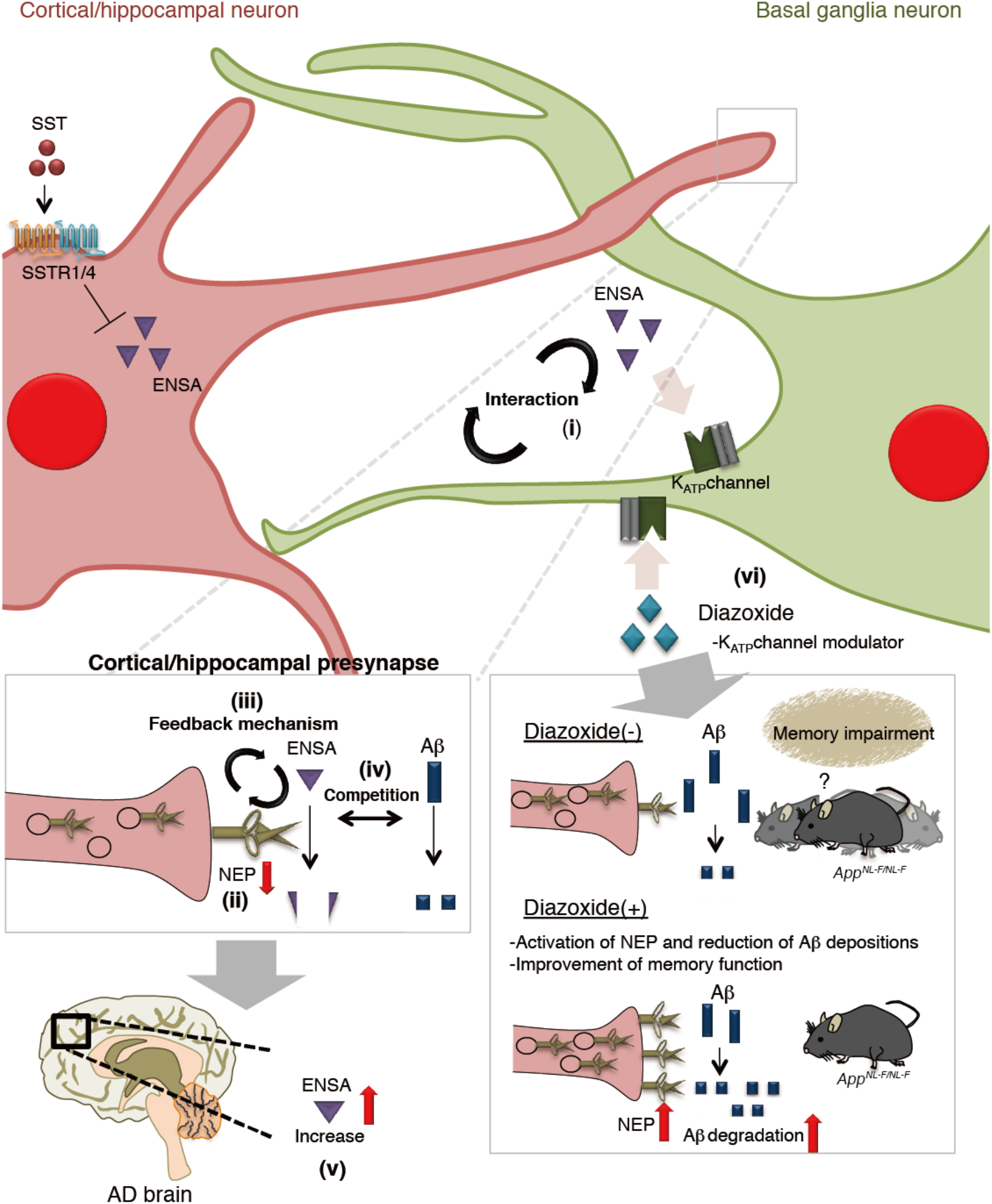
Schematic of key findings of this study. SST-induced NEP activity is regulated by the interaction between cortical/hippocampal and basal ganglia neurons *in vitro* (i). ENSA is a negative NEP regulator (ii). NEP activity may be regulated by a substrate-dependent feedback mechanism (iii). Aβ and ENSA compete with each other in NEP-mediated degradation (iv). ENSA is elevated in aged *App^NL-F^* mice and in patients with AD(iv). K_ATP_ channel modulator diazoxide increases NEP activity and decreases amyloid deposition in *App^NL-F^* mice resulting in improvement of memory function (v).

## Notes

### Competing Interest Statement

The authors have declared no competing interest.

## Reference

1. Hardy, J.A. & Higgins, G.A. Alzheimer’s disease: the amyloid cascade hypothesis. Science 256, 184–185 (1992).

2. Mullan, M., et al. A pathogenic mutation for probable Alzheimer’s disease in the APP gene at the N-terminus of beta-amyloid. Nat Genet 1, 345–347 (1992).

3. Saito, T., et al. Potent amyloidogenicity and pathogenicity of Abeta43. Nat Neurosci 14, 1023–1032 (2011).

4. Rosenberg, R.N., Lambracht-Washington, D., Yu, G. & Xia, W. Genomics of Alzheimer Disease: A Review. JAMA Neurol 73, 867–874 (2016).

5. Iwata, N., et al. Identification of the major Abeta1-42-degrading catabolic pathway in brain parenchyma: suppression leads to biochemical and pathological deposition. Nat Med 6, 143–150 (2000).

6. Iwata, N., et al. Metabolic regulation of brain Abeta by neprilysin. Science 292, 1550–1552 (2001).

7. Yasojima, K., Akiyama, H., McGeer, E.G. & McGeer, P.L. Reduced neprilysin in high plaque areas of Alzheimer brain: a possible relationship to deficient degradation of beta-amyloid peptide. Neurosci Lett 297, 97–100 (2001).

8. Yasojima, K., McGeer, E.G. & McGeer, P.L. Relationship between beta amyloid peptide generating molecules and neprilysin in Alzheimer disease and normal brain. Brain Res 919, 115–121 (2001).

9. Kossut, M., Łukomska, A., Dobrzański, G. & Liguz-Lęcznar, M. Somatostatin receptors in the brain. Postepy Biochem 64, 213–221 (2018).

10. Saito, T., et al. Somatostatin regulates brain amyloid beta peptide Abeta42 through modulation of proteolytic degradation. Nat Med 11, 434–439 (2005).

11. Per Nilsson, K.S., Naomasa Kakiya, Hiroki Sasaguri, Naoto Watamura, Makoto Shimozawa, Satoshi Tsubuki, Risa Takamura, Zhulin Zhou, RaulLoera-Valencia, Misaki Sekiguchi, Aline Petrish, Stefan Schulz, Takashi Saito, Bengt Winblad, Takaomi C. saido. Somatostatin receptor subtypes 1 and 4 redundantly regulate neprilysin, the major amyloid-beta degrading enzyme in brain. bioRxiv (2020). doi:https://doi.org/10.1101/2020.05.09.085795

12. Gunther, T., et al. International Union of Basic and Clinical Pharmacology. CV. Somatostatin Receptors: Structure, Function, Ligands, and New Nomenclature. Pharmacol Rev 70, 763–835 (2018).

13. Heron, L., et al. Human alpha-endosulfine, a possible regulator of sulfonylurea-sensitive KATP channel: molecular cloning, expression and biological properties. Proc Natl Acad Sci U S A 95, 8387–8391 (1998).

14. Virsolvy-Vergine, A., et al. Endosulfine, an endogenous peptidic ligand for the sulfonylurea receptor: purification and partial characterization from ovine brain. Proc Natl Acad Sci U S A 89, 6629–6633 (1992).

15. Saito, T., et al. Single App knock-in mouse models of Alzheimer’s disease. Nat Neurosci 17, 661–663 (2014).

16. Pruitt, A.W., Dayton, P.G. & Patterson, J.H. Disposition of diazoxide in children. Clin Pharmacol Ther 14, 73–82 (1973).

17. Pruitt, A.W., Faraj, B.A. & Dayton, P.G. Metabolism of diazoxide in man and experimental animals. J Pharmacol Exp Ther 188, 248–256 (1974).

18. Zhou, J., et al. Dual sgRNAs facilitate CRISPR/Cas9-mediated mouse genome targeting. Febs j 281, 1717–1725 (2014).

19. Cradick, T.J., Qiu, P., Lee, C.M., Fine, E.J. & Bao, G. COSMID: A Web-based Tool for Identifying and Validating CRISPR/Cas Off-target Sites. Mol Ther Nucleic Acids 3, e214 (2014).

20. Kakiya, N., et al. Cell surface expression of the major amyloid-beta peptide (Abeta)-degrading enzyme, neprilysin, depends on phosphorylation by mitogen-activated protein kinase/extracellular signal-regulated kinase kinase (MEK) and dephosphorylation by protein phosphatase 1a. J Biol Chem 287, 29362–29372 (2012).

21. Rubinfeld, H. & Seger, R. The ERK cascade: a prototype of MAPK signaling. Mol Biotechnol 31, 151–174 (2005).

22. Berndt, N., Dohadwala, M. & Liu, C.W. Constitutively active protein phosphatase 1alpha causes Rb-dependent G1 arrest in human cancer cells. Curr Biol 7, 375–386 (1997).

23. Campbell, D.J. Long-term neprilysin inhibition - implications for ARNIs. Nat Rev Cardiol 14, 171–186 (2017).

24. Solomon, S.D., et al. The angiotensin receptor neprilysin inhibitor LCZ696 in heart failure with preserved ejection fraction: a phase 2 double-blind randomised controlled trial. Lancet 380, 1387–1395 (2012).

25. Skidgel, R.A. & Erdös, E.G. Angiotensin converting enzyme (ACE) and neprilysin hydrolyze neuropeptides: a brief history, the beginning and follow-ups to early studies. Peptides 25, 521–525 (2004).

26. Iwata, N., Higuchi, M. & Saido, T.C. Metabolism of amyloid-beta peptide and Alzheimer’s disease. Pharmacol Ther 108, 129–148 (2005).

27. Roques, B.P., Noble, F., Dauge, V., Fournie-Zaluski, M.C. & Beaumont, A. Neutral endopeptidase 24.11: structure, inhibition, and experimental and clinical pharmacology. Pharmacol Rev 45, 87–146 (1993).

28. Turner, A.J., Isaac, R.E. & Coates, D. The neprilysin (NEP) family of zinc metalloendopeptidases: genomics and function. Bioessays 23, 261–269 (2001).

29. Johnson, G.D., Stevenson, T. & Ahn, K. Hydrolysis of peptide hormones by endothelin-converting enzyme-1. A comparison with neprilysin. J Biol Chem 274, 4053–4058 (1999).

30. Hama, E., Shirotani, K., Iwata, N. & Saido, T.C. Effects of neprilysin chimeric proteins targeted to subcellular compartments on amyloid beta peptide clearance in primary neurons. J Biol Chem 279, 30259–30264 (2004).

31. Tsubuki, S., Takaki, Y. & Saido, T.C. Dutch, Flemish, Italian, and Arctic mutations of APP and resistance of Abeta to physiologically relevant proteolytic degradation. Lancet 361, 1957–1958 (2003).

32. Miyakawa, T., et al. Neurogranin null mutant mice display performance deficits on spatial learning tasks with anxiety related components. Hippocampus 11, 763–775 (2001).

33. Alvarez-Fernandez, M., et al. Greatwall is essential to prevent mitotic collapse after nuclear envelope breakdown in mammals. Proc Natl Acad Sci U S A 110, 17374–17379 (2013).

34. Rangone, H., et al. Suppression of scant identifies Endos as a substrate of greatwall kinase and a negative regulator of protein phosphatase 2A in mitosis. PLoS Genet 7, e1002225 (2011).

35. Burgess, A., et al. Loss of human Greatwall results in G2 arrest and multiple mitotic defects due to deregulation of the cyclin B-Cdc2/PP2A balance. Proc Natl Acad Sci U S A 107, 12564–12569 (2010).

36. Gahete, M.D., et al. Expression of Somatostatin, cortistatin, and their receptors, as well as dopamine receptors, but not of neprilysin, are reduced in the temporal lobe of Alzheimer’s disease patients. J Alzheimers Dis 20, 465–475 (2010).

37. Saito, T., Takaki, Y., Iwata, N., Trojanowski, J. & Saido, T.C. Alzheimer’s disease, neuropeptides, neuropeptidase, and amyloid-beta peptide metabolism. Sci Aging Knowledge Environ 2003, Pe1 (2003).

38. Saido, T.C. & Iwata, N. Metabolism of amyloid beta peptide and pathogenesis of Alzheimer’s disease. Towards presymptomatic diagnosis, prevention and therapy. Neurosci Res 54, 235–253 (2006).

39. Davies, P., Katzman, R. & Terry, R.D. Reduced somatostatin-like immunoreactivity in cerebral cortex from cases of Alzheimer disease and Alzheimer senile dementa. Nature 288, 279–280 (1980).

40. Lu, T., et al. Gene regulation and DNA damage in the ageing human brain. Nature 429, 883–891 (2004).

41. Wang, T.L., Chang, H., Hung, C.R. & Tseng, Y.Z. Morphine preconditioning attenuates neutrophil activation in rat models of myocardial infarction. Cardiovasc Res 40, 557–563 (1998).

42. Joshi, D.D., et al. Negative feedback on the effects of stem cell factor on hematopoiesis is partly mediated through neutral endopeptidase activity on substance P: a combined functional and proteomic study. Blood 98, 2697–2706 (2001).

43. Liu, D., et al. The KATP channel activator diazoxide ameliorates amyloid-beta and tau pathologies and improves memory in the 3xTgAD mouse model of Alzheimer’s disease. J Alzheimers Dis 22, 443–457 (2010).

44. Oddo, S., Caccamo, A., Kitazawa, M., Tseng, B.P. & LaFerla, F.M. Amyloid deposition precedes tangle formation in a triple transgenic model of Alzheimer’s disease. Neurobiol Aging 24, 1063–1070 (2003).

45. Ott, T. & Nieder, A. Dopamine and Cognitive Control in Prefrontal Cortex. Trends Cogn Sci 23, 213–234 (2019).

46. Wu, Y.N., Shen, K.Z. & Johnson, S.W. Differential actions of AMP kinase on ATP-sensitive K(+) currents in ventral tegmental area and substantia nigra zona compacta neurons. Eur J Neurosci 46, 2746–2753 (2017).

47. Knowlton, C., Kutterer, S., Roeper, J. & Canavier, C.C. Calcium dynamics control K-ATP channel-mediated bursting in substantia nigra dopamine neurons: a combined experimental and modeling study. J Neurophysiol 119, 84–95 (2018).

48. Schiemann, J., et al. K-ATP channels in dopamine substantia nigra neurons control bursting and novelty-induced exploration. Nat Neurosci 15, 1272–1280 (2012).

49. Gasbarri, A., Packard, M.G., Campana, E. & Pacitti, C. Anterograde and retrograde tracing of projections from the ventral tegmental area to the hippocampal formation in the rat. Brain Res Bull 33, 445–452 (1994).

50. Lisman, J.E. & Grace, A.A. The hippocampal-VTA loop: controlling the entry of information into long-term memory. Neuron 46, 703–713 (2005).

51. Yokoshiki, H., Sunagawa, M., Seki, T. & Sperelakis, N. ATP-sensitive K+ channels in pancreatic, cardiac, and vascular smooth muscle cells. Am J Physiol 274, C25–37 (1998).

52. Guan, L., et al. Diazoxide induces endoplasmic reticulum stress-related neuroprotection mediated by p38 MAPK against Abeta25-35 insults. Eur Rev Med Pharmacol Sci 22, 6133–6138 (2018).

53. Kong, M. & Ba, M. Protective effects of diazoxide against Abeta(2)(5)(-)(3)(5)-induced PC12 cell apoptosis due to prevention of endoplasmic reticulum stress. Neuroreport 23, 493–497 (2012).

54. Tan, S., et al. Effects of Abeta1-42 on the current of KATP channels in cultured cholinergic neurons. Neurol Res 34, 707–713 (2012).

55. Fu, Q., et al. Diazoxide pretreatment prevents Abeta1-42 induced oxidative stress in cholinergic neurons via alleviating NOX2 expression. Neurochem Res 39, 1313–1321 (2014).

56. Virgili, N., et al. K(ATP) channel opener diazoxide prevents neurodegeneration: a new mechanism of action via antioxidative pathway activation. PLoS One 8, e75189 (2013).

57. Lu, B., et al. Neutral endopeptidase modulation of septic shock. J Exp Med 181, 2271–2275 (1995).

58. Wiśniewski, J.R., Zougman, A., Nagaraj, N. & Mann, M. Universal sample preparation method for proteome analysis. Nat Methods 6, 359–362 (2009).

59. Fujii, W., Kawasaki, K., Sugiura, K. & Naito, K. Efficient generation of large-scale genome-modified mice using gRNA and CAS9 endonuclease. Nucleic Acids Res 41, e187 (2013).

60. Hsu, P.D., et al. DNA targeting specificity of RNA-guided Cas9 nucleases. Nat Biotechnol 31, 827–832 (2013).

